# Beyondcell: targeting cancer therapeutic heterogeneity in single-cell RNA-seq

**DOI:** 10.1101/2021.04.08.438954

**Authors:** Coral Fustero-Torre, María José Jiménez-Santos, Santiago García-Martín, Carlos Carretero-Puche, Luis García-Jimeno, Tomás Di Domenico, Gonzalo Gómez-López, Fátima Al-Shahrour

## Abstract

We present Beyondcell (https://gitlab.com/bu_cnio/beyondcell/), a computational methodology for identifying tumour cell subpopulations with distinct drug responses in single-cell RNA-seq data and proposing cancer-specific treatments. Our method calculates an enrichment score in a collection of drug signatures, delineating therapeutic clusters (TCs) within cellular populations. Additionally, Beyondcell determines therapeutic differences among cell populations, and generates a prioritised ranking of the differential sensitivity drugs between chosen conditions to guide drug selection. We performed Beyondcell analysis in four single-cell datasets to validate our score and to demonstrate that TCs can be exploited to target malignant cells both in cancer cell lines and tumour patients.

## Background

Tumour heterogeneity (TH) refers to genetic and epigenetic differences of the same tumour type between patients, between tumours in a single patient, and within the cells of a tumour. It contributes to the medical complexity of cancer treatment and can lead to drug resistance, therapeutic failure and higher lethal outcome (1). TH is closely related to tumour evolution which has been described as branching clonal evolution due to the accumulation of somatic mutations (2). The heterogeneity of cancer cells, tumour evolution and clonal dynamics introduce significant challenges in designing effective treatment strategies. Unfortunately, these issues are not satisfactorily addressed in routine clinical practice as systematic efforts to consider inter-patient TH when choosing therapies are still limited, while they are extremely rare for intra-TH (3, 4)

Single-cell RNA sequencing (scRNA-seq) has become an established technology to dissect TH at the transcriptional level, revealing high-resolution cellular composition and dynamics and offering an unprecedented opportunity to address TH therapeutically (5, 6, 7). For instance, scRNA-seq studies have been successfully employed to identify novel cancer cell subpopulations (8), tumour biomarkers (9), drug resistance pathways and therapeutic targets (10). In addition, large-scale projects focused on obtaining a comprehensive molecular and pharmacological characterisation of the cancer cell lines have provided valuable datasets relating gene expression signatures with drug response and treatment sensitivity. Some examples include the Cancer Cell Line Encyclopedia (CCLE) (11), the Genomics of Drugs Sensitivity in Cancer (GDSC) (12), the Cancer Therapeutic Response Portal (CTRP) (13) and the Drug Repurposing Hub/LINCS (14).

In this situation, it is reasonable to hypothesise that drugs (or drug combinations) capable of targeting TH at single-cell resolution can be identified by integrating drug response profiles and scRNA-seq data. This would help to address the tumour therapeutic complexity, revealing the impact of TH in response to drugs and the scope of tumour cells whose therapeutic approach could be managed with approved drugs, clinical trials or drug repositioning strategies. However, there are no current computational methodologies capable of relating single-cell gene expression and high-throughput drug screening datasets to suggest knowledge-driven treatments. To address these challenges we developed Beyondcell, a novel method for detecting tumour cell subpopulations with distinct drug response in order to estimate the tumour therapeutic complexity and propose differential drugs to target cell subpopulations, experimental conditions or phenotypes in scRNA-seq experiments. We have applied Beyondcell to four different studies, including both cell lines and patient-based single-cell data. Our analysis covers the detection of TCs, as well as the visualization of results for clinical interpretation.

## Methods

### Beyondcell workflow and therapeutic clusters

An analysis with Beyondcell starts with a single-cell expression matrix and a collection of drug signatures: the drug perturbation (PS_C_) and the drug sensitivity (SS_C_) collections containing 4690 and 819 signatures respectively (11, 12, 13, 14). PS_C_ captures the transcriptional changes induced by a drug, while SS_C_ contains signatures reflecting the transcriptional status of sensitivity or resistance prior to drug treatment (**Fig. 1a**).

**Figure 1.**
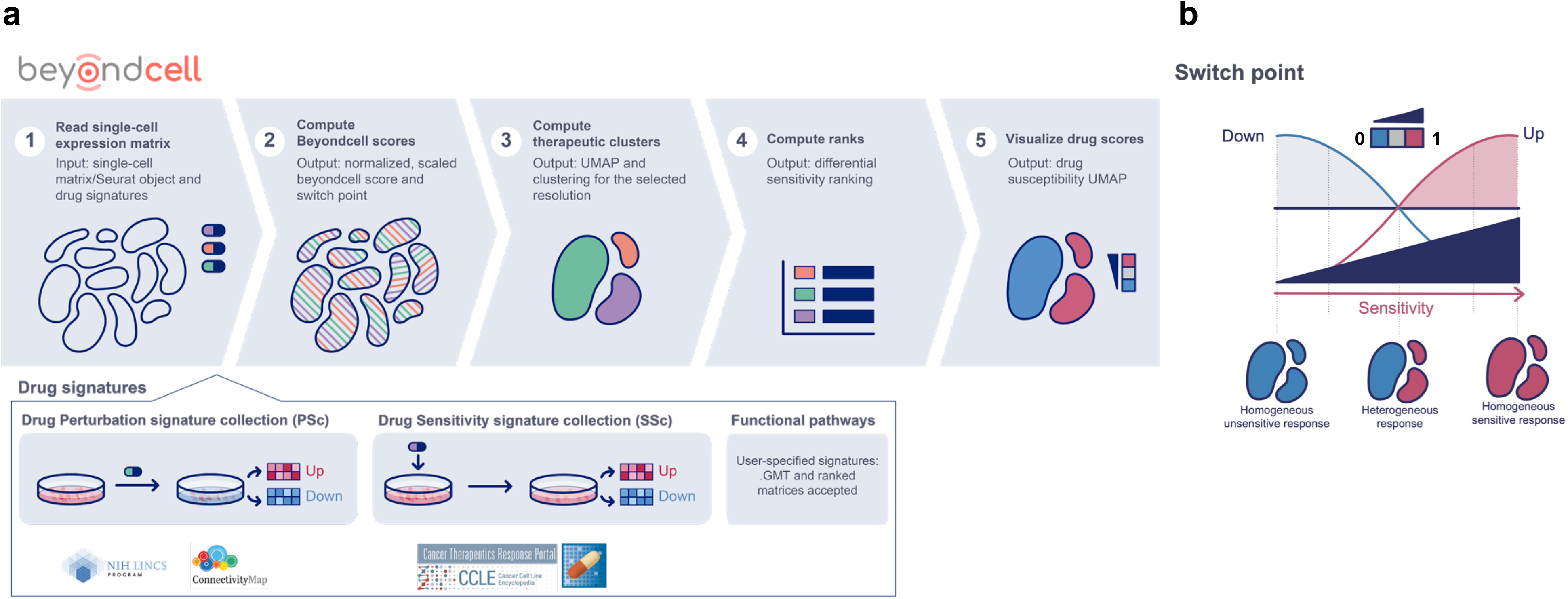
The Beyondcell workflow and Beyondell switch point. Beyondcell is a methodology for the identification of drug vulnerabilities in scRNA-seq data. Using Beyondcell, we have identified the presence of therapeutic clusters in our data, defined as sets of cells sharing a common behaviour towards a collection of drugs. **(a)** Given two inputs, an scRNA-seq expression matrix and a drug signature collection — either the drug perturbation (PS_C_) or the drug sensitivity (SS_C_) collections or a user-provided GMT file/ranked matrix — Beyondcell calculates a score (BCS) for each drug-cell pair. The resulting BCS matrix is used to determine the presence of therapeutic clusters, which can be visualised using a UMAP in Beyondcell. A sensitivity-based ranking can be obtained in order to prioritise the best hits. (**b)** The scaled BCS ranges from 0 to 1 and measures the cell perturbation susceptibility (when using the PS_C_) or the predicted sensitivity to a given drug (when using the SS_C_). The BCS can also be used to evaluate the cells’ functional status if functional signatures are applied. Furthermore, a switch point (SP) is calculated for each analysed signature, by determining the value in the 0 to 1 scale where cells switch from a down-regulated status to an up-regulated one. Thus, the most therapeutically-homogeneous tumours would be those in which each and every one of their cells responds with the same directionality to a certain drug, either towards sensitivity (SP = 0) or resistance (SP = 1), while a heterogeneous response would be represented by intermediate SPs.

Our method calculates the Beyondcell Score (BCS) which estimates, for each cell in the preprocessed single-cell expression matrix, the enrichment in every drug signature in the specified collections. The BCS ranges from 0 to 1 and measures the cell perturbation susceptibility (using the PS_C_) or the predicted sensitivity to a given drug (using the SS_C_). The BCS can also evaluate the cells’ functional status using functional gene sets such as molecular pathways, or cancer hallmarks. The calculated BCS matrix allows the determination of therapeutic clusters (TCs) within cellular populations defined as ‘a set of cells that share a common response to a set of drugs’. In order to find potential TCs, a clustering analysis is applied to the BCS matrix where cells are grouped by their differential response to the selected drugs. For each drug, Beyondcell also calculates the switch point (SP), which reflects the homogeneity of the drug response throughout the single-cell dataset **(Fig. 1b)**. Thus, the most therapeutically homogeneous tumours would be those in which each and every one of their cells responds in the same way to a certain drug, either with a sensitivity (SP=0) or resistance (SP=1) response, while a heterogeneous response would be represented by intermediate SPs. Then, the BCS matrix and the SP are used to generate a Uniform Manifold Approximation and Projection (UMAP), enabling the visualisation of the TCs reflecting the homogeneity in the drug responses within cellular populations, and to highlight in TCs drug response, experimental conditions, cell subtypes, biomarker expression and cellular functional activity. Additionally, Beyondcell determines therapeutic differences among cell populations and generates a prioritised ranking of the differential sensitivity drugs between chosen conditions to guide drug selection (Supplementary Methods).

### Data description

The Library of Integrated Network-based Cellular Signatures (LINCS, http://www.lincsproject.org) (14) is a catalogue of gene expression data associated with cell lines exposed to a variety of perturbing agents, such as small molecules (about 5500), FDA drugs (~1300) and shRNA silencing (22,119 genetic perturbagens). LINCS, based on L1000 high throughput technology, is an extension of the Connectivity Map, which has been successfully used for drug repositioning (15). From the ~20K small molecules tested in the LINCS L1000 dataset, only those with an identifiable common drug name were selected. This reduced the number of drugs to 4690. In the CCLE (11) 28 drugs were tested and the mRNA expression of 1037 cell lines were profiled using Affymetrix U133Plus2 arrays. The gene-centric RMA-normalised mRNA expression data, the cell line information and the drug response data from Cancer Cell Line Encyclopedia were downloaded from the CCLE data portal (https://portals.broadinstitute.org/ccle/home). In the CTRP (13), 545 drugs (single or combination) were tested against 887 cell lines. We downloaded the drug response data of the CRTP 2.1 project from the CTD2 Data Portal (https://ocg.cancer.gov/programs/ctd2/data-portal). The gene expression data used in order to obtain the CTRP expression signatures were those of the CCLE. Of the 887 cell lines, 805 were present in the CCLE dataset. In the GDSC data (12), 265 compounds corresponding to 250 different drugs were tested against 1074 cell lines. Cell lines were profiled for gene expression using the Affymetrix Human Genome U219 array. Expression data were normalised using RMA (16). Drug response data and preprocessed mRNA gene expression data were downloaded from the GDSC web portal (http://www.cancerrxgene.org/downloads).

### Signature generation

A gene expression signature is a general model for the representation of the transcriptional changes associated to a given biological phenotype or perturbation. In this study we have considered two expression signature collections: the PS_C_ captures the transcriptional changes induced by a drug; while the SS_C_ captures the sensitivity to the drug effect. In both cases, these expression signatures are obtained from a differential expression analysis, although using different designs. Despite its different biological interpretations, both types of signature were represented as the two gene sets formed by the *N* most up and down-regulated genes (the *UP* and *DN* genesets). Several *N* were tested (50, 100, 250 and 500) and no significant disagreement was found between the top synergistic and antagonistic interactions (data not shown). Thus, following (17), *N* was taken as 250.

### Drug perturbation signature collection (PS_C_)

Drug-induced expression signatures were obtained from experiments in which the transcriptional state of the cell is measured before and after treatment with the drug. This makes it possible to study the transcriptional effect of the drug. In order to obtain consensus expression signatures for each drug, a differential expression analysis was performed on control vs treated cells using *limma* (18). Full details of PS_C_ signature collection are available at Supplementary Methods.

### Drug sensitivity signature collection (SS_C_)

Drug-induced expression signatures were obtained from pharmacogenomics experiments in which the pharmacological response to a drug and the transcriptional state before treatment were considered. In order to obtain the expression signature, a differential expression analysis against a measure of drug response was performed using *limma.* The area under the curve (AUC) was used as the measure of drug response, as contrary to IC_50_, it can always be estimated without extrapolation from the dose-response curve, and also because it has shown more accuracy in the prediction of drug response (19). In all the projects considered, the tumour origin of the cell lines were considered as confounding variables. In the CCLE the covariate was created by combining the *Site.Primary* and the *Histology* information of the cell line. In the GDSC 2.0, the variable *Site* and in the CTRP, the ‘*CCLE primary Site*’ was used. Further details of SS_C_ signature collection are available at Supplementary Methods.

### Functional Pathways

The Beyondcell package also includes a small collection of curated functional pathways obtained from the Molecular Signatures Database (MSigDB). These are meant to give the user some insight about the cells’ viability and status. These pathways are related to the regulation of the epithelial-mesenchymal transition (EMT), cell cycle, proliferation, senescence and apoptosis. Furthermore, the package is able to accept external signatures in GMT format or as ranked matrices.

### Beyondcell score calculation

The Beyondcell score evaluates the activity of a signature of interest in a single-cell RNA-seq experiment. The transcriptomics data needs to be pre-processed, meaning that proper cell-based quality control filters, as well as normalisation, scaling and clustering of the data, should be applied prior to the analysis with Beyondcell. When analysing a bidirectional gene signature (a signature with separate sets of up-regulated and down-regulated genes), the Beyondcell score is independently obtained for each signature mode. The individual sum of the expression is calculated and divided by the number of genes in the given signature that are present in the single-cell expression matrix. The obtained raw scores are normalised and the individual up and down normalised scores are summed and rescaled between 0 and 1. In cases where the gene signature is unidirectional, all steps remain the same, although the rescaling will only be applied to one set of normalised scores.

### Drug-background selection

In order to obtain TCs, the Beyondcell methodology can optionally generate a background score matrix. The background score matrix allows the user to reduce the computation time. It aims to characterise the heterogeneity of the drug responses in the whole dataset, by using a reduced collection of drugs that is able to capture the main differences between individual cells. To generate this background selection, the drug specificity score (DSS) of the whole PS_C_ collection has been calculated and the first and last decile have been selected. Here, the rationale is the following: for each drug, the DSS score quantifies the similarity between the induced expression patterns across cell types. And while some drugs induce similar patterns across distinct cell types, the majority of them have different effects across all of them (20).

### Regression of unwanted sources of variation

After obtaining the normalised and scaled Beyondcell scores, we observed that the normalisation step was not sufficient to avoid the correlation between sample-level metrics (number of genes, number of UMIs or even cell cycle status) and the calculated scores. To correct for this, we have implemented a regression function. *bcRegressOut* removes the effect of all unwanted sources of variation from the scoring matrix in two steps: first, it imputes missing data (if necessary) using a k-Nearest Neighbours (KNN) algorithm; second, it obtains the residuals derived from the regression model via QR decomposition. The obtained residuals can then be used for the downstream dimensional reduction steps.

### Therapeutic clusters

The BCS matrix, including the computed scores for all drugs of interest, can be used as an input for a dimensionality reduction and clustering analysis. In this step, cells are grouped by their differential response to the analysed drugs. A Uniform Manifold Approximation and Projection (UMAP) allows the visualisation of the identified TCs. With this analysis, cells can be classified into distinct TCs, which represent sets of cells sharing a common response to a particular drug exposure.

### Drug prioritisation

The *bcRank* function computes the BCS matrix statistics. In particular, it calculates the switch point, mean, median, standard deviation, variance, minimum, maximum, proportion of NaN and residuals’ mean of each signature. We recommend prioritising drugs taking into account both the switch point and the residuals. The Beyondcell package includes the function *bc4Squares,* which helps to visualise this prioritisation. A 4 squares plot consists in a scatter plot of the residuals’ means (x axis) vs the switch points (y axis) of a specified cluster (either a TC or a group defined by experimental condition or phenotype). The 4 quadrants are highlighted: the top left and bottom right corners contain the drugs to which these cells are least/most sensitive, respectively. The centre quadrants show the drugs to which the selected cells are differentially insensitive or sensitive when compared to the other clusters.

## Results

### Reliability of the Beyondcell score and its application in cancer cell lines under drug exposure

We applied Beyondcell to the Ben-David *et al*. dataset (21) to demonstrate the reliability of the BCS and to validate its usefulness for identifying drug-response cell subpopulations. This study dissects the genetic and transcriptional heterogeneity within cancer cell lines, providing an scRNA-seq dataset that includes 7101 cells obtained from one single-cell-derived clone from the MCF7 cell line (MCF7-AA) exposed to bortezomib. Cells were collected before and after 12, 48 and 96 hours of drug exposure (t0, t12, t48 and t96) to study its antiproliferative effects. Also, a drug screening of 321 anticancer compounds was performed to study drug response heterogeneity across 27 strains of the MCF7 breast cancer cell line.

First, to demonstrate the reliability of the BCS, we computed BCS for each compound using SS_C_ drug signatures to the collected MCF7-AA cells at t0. We found that median BCS obtained with SS_C_ signatures for cells at t0 correlate significantly (R = −0.26, p < 1e−04) with MCF7-AA cell viability measures reported by Ben-David *et al.* after treatment, demonstrating that BCS reflects drug sensitivity (**Fig. 2a).** When we employed PS_C_ signatures, drugs were then classified in three groups: chemotherapy, targeted therapy and others (including immunotherapy, hormone therapy and photodynamic therapy). The BCS for targeted therapies showed a significant correlation with MCF7-AA cell viability (R = −0.24, p = 8.4e−03) (Additional File 1: **Fig. S1A**) while the rest of the therapies were not found significant. This could suggest that PS_C_ signatures for targeted therapies reflect more accurately which cells are more likely to respond than chemotherapy signatures.

**Figure 2.**
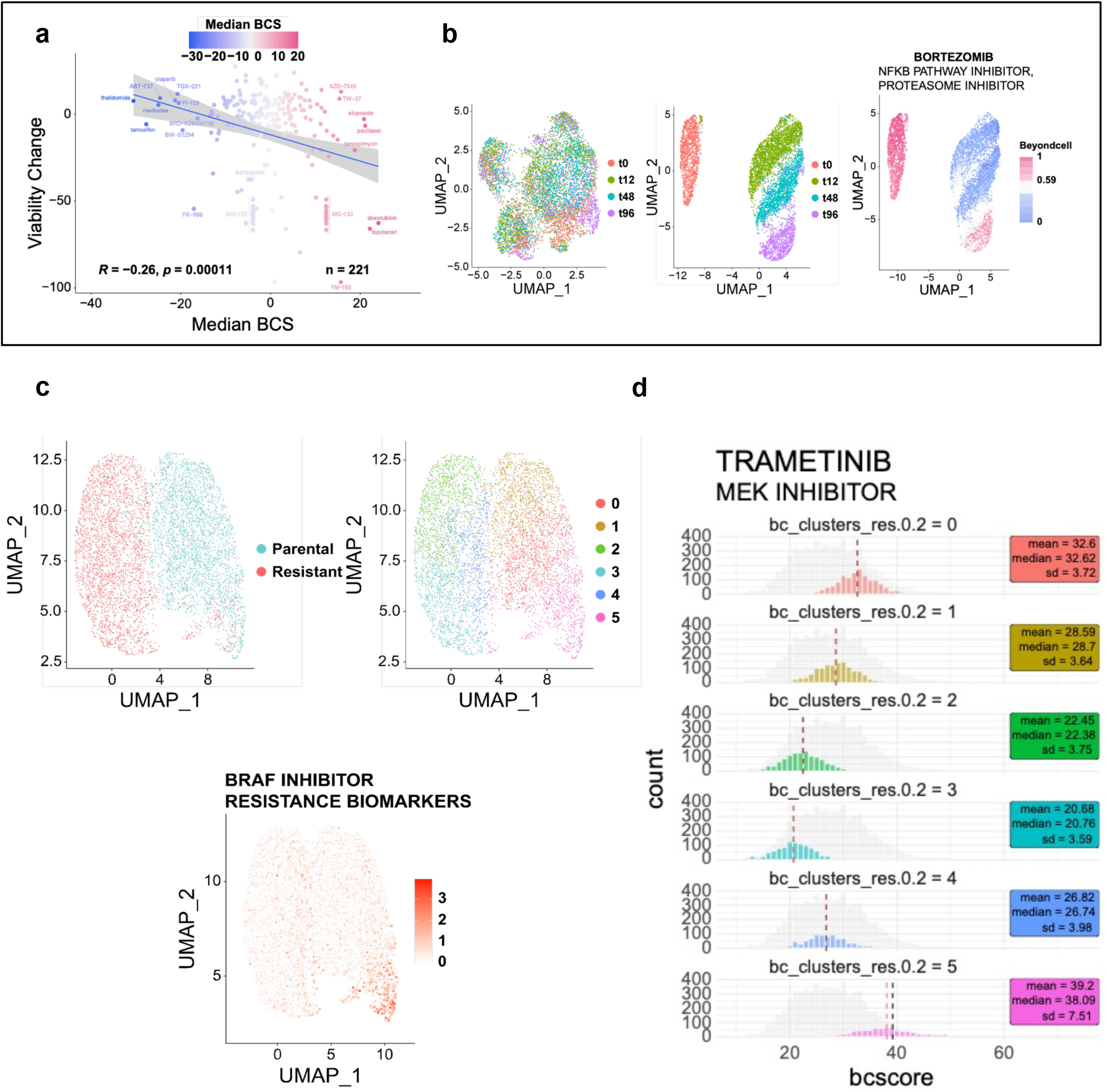
Reliability of the Beyondcell score and its application in cancer cell lines under drug exposure. **(a)** Median BCS obtained with SS_C_ signatures for cells at t0 correlate negatively (R = −0.26, p < 1e−04) with MCF7-AA viability measures reported by Ben-David *et al*. **(b)** Using BCS obtained with PS_C_ signatures, Beyondcell is capable of clustering MCF7-AA cells treated with bortezomib (Ben-David *et al.*) into therapeutic clusters that overlap with treatment times. *Left:* UMAP plot of the integrated Seurat object coloured by treatment time; *Centre:* UMAP plot of the Beyondcell object also coloured by treatment time; *Right:* UMAP plot of the Beyondcell object coloured by bortezomib BCS. Untreated cells (t0) are sensitive to bortezomib whereas cells undergoing treatment (t12 and t48) are insensitive. After treatment during 72h followed by drug wash and 24h of recovery, cells at 96h (t96) restore their sensitivity to this drug. **(c)** Beyondcell UMAP plot for the Ho *et al.* dataset and the SS_C_ drug signatures. *Top left:* cells coloured according to the treatment condition; *Top right:* cells coloured by Beyondcell’s therapeutic clusters; *Bottom:* Beyondcell UMAP plot showing the summed expression of several BRAF inhibitor-resistant biomarkers (*JUN*, *WNT5A*, *PDGFRB*, *EGFR*, *NRG1*, *FGFR1* and *AXL*). **(d)** Histogram representing trametinib Beyondcell scores in each therapeutic cluster.

Next, we were interested in evaluating Beyondcell’s ability to identify distinct drug-response cell subpopulations before and after bortezomib exposure. Beyondcell’s analysis with the PS_C_ collection clearly separates the cells into discrete clusters, in contrast to the mixed cell groups found using transcriptional profiles. The resulting therapeutic clusters not only separated bortezomib-treated and untreated cells but also retrieved drug exposure time-points. By focusing on a PS_C_ bortezomib expression signature, t12 and t48 cells reflected the perturbation status induced by bortezomib in contrast to t0 cells. Interestingly, t96 cells reverted to a pre-perturbation status for the bortezomib signature, highlighting the reversible inhibitory capacity of this proteasome inhibitor (22) (**Fig. 2b)**. In particular, cells were more sensitive to bortezomib when combined with SNX-2112, an *HSP90* inhibitor, than in treatment with bortezomib alone, suggesting its role as a sensitiser to bortezomib treatment (Additional File 1: **Fig. S1B**). In summary, in this study we validate the BCS as a measure of drug sensitivity, determining its significant correlation with drug screening experimental data. In addition, we demonstrate Beyondcell’s utility to recapitulate drug response in the MCF7 cancer cell line and propose single and drug combinations to sensitise the cells.

### Analysis of BRAF inhibitor sensitive and resistant subpopulations in melanoma cells

Next, using data from a 451HLu human melanoma cell line treated with BRAF inhibitors (BRAFi), we evaluated Beyondcell’s ability to identify drug-resistant cellular populations and to propose drug treatments (23). Beyondcell analysis with SS_C_ drug signatures recapitulated ‘parental’ and ‘BRAFi*-*resistant’ cell populations and identified 6 different therapeutic clusters shown in a Beyondcell UMAP plot where cells are coloured according to the treatment condition or the Beyondcell therapeutic cluster. Therapeutic cluster 5 (TC5) included both parental and resistant cells. A small fraction of the parental cells in TC5 (15%) expressed BRAFi-resistance markers like *AXL*, *NRG1*, *DCT* and *FGFR1* that are also expressed in BRAFi-resistant cells, showing that they were clonally selected from the parental population contributing to the resistance (**Fig. 2c;** Additional File 1: **Fig. S2**).

Additionally, Beyondcell identified specific drugs to target TC5, BRAFi-resistant and parental cells. For instance, cells in cluster 5 are differentially sensitive to dasatinib (SRCi) and unresponsive to dinaciclib (CDKi) and gemcitabine. On the other hand, cells in the resistant condition are differentially sensitive to bardoxolone methyl (NF-kBi) and unresponsive to AZD6482 (PIK3i) (Additional File 1: **Fig. S3; Table S1**). MEK inhibitors (MEKi) are shown to target 451HLu cells including pre-resistant clones (TC5). For instance, trametinib had positive Beyondcell scores in each therapeutic cluster obtained for SS_C_ drug signatures, showing a higher BCS score (higher sensitivity) in TC5, and therefore, it could be proposed to target BRAFi-unresponsive cells. In fact, MEKi is a standard treatment in advanced melanoma patients in combination with BRAFi (24) (**Fig. 2d;** Additional File 1: **Fig. S4A**). We observed that TC5 cells were also grouped in expression UMAPs overlapping with the scRNA-seq expression cluster 2 (EC2), suggesting a direct relationship between the drug response profiles and gene expression patterns (Additional File 1: **Fig. S4B)**. In order to identify novel genes involved in BRAFi-resistance, we performed a differential gene expression analysis for TC5, revealing 572 significantly overexpressed genes (|log_2_(FC)| > 2, FDR < 0.05) (**Table S2),** including some known BRAFi-resistance biomarkers (e.g. *AXL* and *NRG1*). In addition, vemurafenib and dabrafenib BCSs obtained using SS_C_ showed that TC5 had lower sensitivity to RAF inhibitors than the rest of the TCs, confirming that TC5 cells are pre-resistant to BRAFi (Additional File 1: **Fig. S4C).** This validation demonstrates that Beyondcell is able to identify therapeutic clusters that recapitulate drug effects on cells, propose drugs to target sensitive and resistant cells, and identify drug-response biomarkers.

### Beyondcell characterises single-cell variability in drug response in pan-cancer cell line data

We also applied Beyondcell to describe the therapeutic heterogeneity in 198 cell lines from 22 cancer types (25). Using SS_C_ drug signatures we found 5 TCs where 12 of 22 cancer types were overrepresented in at least one of these TCs. TC0 was mostly constituted by cells from skin cancer (melanoma) and endometrial/uterine cancer cell lines, while TC1 included cells from bladder, gallbladder and pancreatic cancer cell lines. TC3 was enriched in breast and colon/colorectal cancer, while gliomas were exclusively located in TC2 together with kidney and thyroid cancer (Additional File 1: **Fig. S5; Table S3)**. TC4 was mainly constituted by two cell lines: the osteosarcoma (HOS) and sarcoma (A204) cell lines. Interestingly, 11 out of 12 brain cancer cell lines clustered together in TC2 independently of whether the lineage is astrocytoma or glioblastoma, with the exception of the single medulloblastoma cell line that is located in TC0. These results suggest that the cell lines from these 12 cancer types that tend to cluster in the same TCs have a common drug response.

In contrast, cancer types such as lung, gastric and ovarian among others showed high cellular therapeutic heterogeneity. For instance, the four bile duct cancer cell lines are each grouped in a different TC showing different drug response profiles despite belonging to the same cancer type (Additional File 1: **Fig. S6A)**. Interestingly, lung cancer cell lines are distributed between the TCs regardless of whether they are squamous or adenocarcinoma subtype showing diverse drug response (Additional File 1: **Fig. S6B**). We also tested whether these lung cancer cell lines expressing this distinct drug response pattern exhibit a unique pattern of mutations and genetic vulnerabilities. For this, we used the CCLE mutation dataset as well as the Achilles dataset of genome-wide CRISPR knockout screens to map known driver mutations in lung cancer (e.g. *KRAS*, *EGFR*, *PI3KCA*) and the genes identified as essential for proliferation (26). These analyses showed that lung cancer cell lines did not cluster together, therefore, TCs detected by Beyondcell are not driven by mutational and essentiality events (Additional File 1: **Fig. S6C**).

We hypothesised that the heterogeneous therapeutic landscape observed in cancer cell lines could also be an opportunity to infer drug repositioning strategies to target cell lines with different tissue origin or genetic background but clustered together in Beyondcell by similar drug responses (27). For instance, most of the colon/colorectal cancer cell lines were clustered in TC3 except cells from the NCIH747 colorectal cell line, which are mostly concentrated in TC1 where bladder, gallbladder and pancreatic cancer cell lines are located (Additional File 1: **Fig. S7**). This suggests that NCIH747 cell line could respond to tyrosine kinase inhibitors (TKIs) prescribed by Beyondcell for these tumour types such as EGFRi and also to inhibitors to target members from MAPK pathway (MEK and SRC). Interestingly, NCIH747 cell line has shown experimental sensitive response to selumetinib, a MEKi, this drug being the most differential sensitive drug for TC1 and for bladder and pancreatic cancer in our results. These findings highlight Beyondcell’s ability to propose drugs for repurposing, providing additional drug response information using transcriptional data that could complement diagnostic information (i.e. tissue origin, stage, mutational status, etc) that commonly guide cancer treatment.

Overall, global expression profiles clustered cells by cancer type, however, the TCs did not show such separation, suggesting less variability in the drug response. More specifically, 172 of 198 cell lines were overrepresented in the same TC, indicating that these cell lines had a lower cellular therapeutic heterogeneity than the rest of the cell lines (n = 26), which were spread across the different TCs (**Table S3)**. The high-grade serous ovarian cancer cell lines are a clear example of high therapeutic heterogeneity, since cells from these cell lines are mostly distributed between TCs (Additional File 1: **Fig. S8**). This observed differential drug sensitivity led by varying patterns of gene expression could result from clonal dynamics and continuous genetic instability that translates into heterogeneity in cancer cell lines (21).

Beyondcell results for SS_C_ revealed 136 differential sensitivity drugs for TCs and 183 drugs in cancer type comparison. In general, TC1 and TC4 showed higher sensitivity to EGFR and topoisomerase inhibitors respectively while TC0 and TC3 both showed higher sensitivity to PLK and CDK inhibitors (**Fig. 3a;** Additional File 1: **Fig. S9).** Using PS_C_ 174 drugs showed differential sensitivity for TCs and 569 drugs in cancer type comparison **(Table S4).** However, TCs did not form discrete clusters so we expect that changes in drug responses are subtle, with a lot of commonalities. Beyondcell was also used to compute drug-response similarity correlation modules using BCS matrix (Additional File 1: **Fig. S10; Table S5).** These correlation modules could be used to infer therapeutic mechanisms of action (MoA) for those drugs with unknown targets.

**Figure 3.**
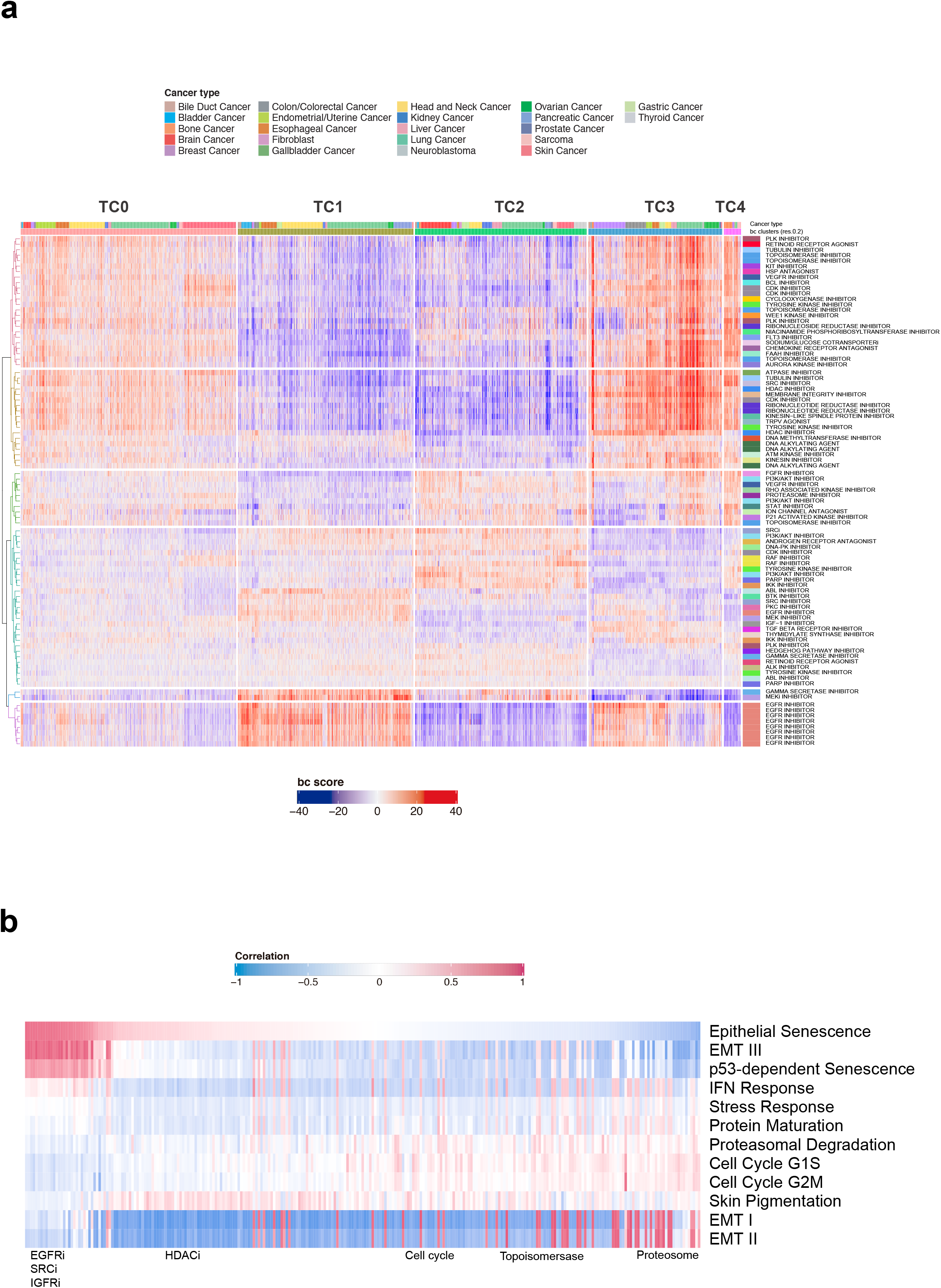
Beyondcell characterises single-cell variability in drug response in pan-cancer cell line data. **(a)** The heatmap depicts both the cluster and cancer-specific drugs with a heterogeneous sensitivity pattern found in Kinker *et al*. Only drugs with a mechanism-of-action (MoA) annotation are shown. **(b)** We found EGFR inhibitors to be highly correlated with the Epithelial Senescence, EMT III and p53-dependent senescence programs, while Tyrosine Kinase inhibitors or Topoisomerase inhibitors were anticorrelated.

Beyondcell predicts EGFRi as the most enriched drugs for TC1, but also, for bladder and gallbladder cancer. To validate this result, we used a recently generated dataset of clinical compounds screened across 578 cell lines (27). Differential drug vulnerability analysis showed increased sensitivity to EGFR inhibitors (Additional File 1: **Fig. S11; Table S6**) for cell lines in TC1, while TC2 showed decreased sensitivity, confirming our results. Moreover, *EGFR* is overexpressed in up to 74% of bladder cancer tissue specimens but has a relatively low expression in normal urothelium suggesting that it could be a potential therapeutic target. In addition, *EGFR* is an independent predictor of decreased survival and stage progression in bladder cancer (28).

Beyondcell calculates an SP for every drug, providing a measure of drug response homogeneity and sensitivity in each cell line. For instance, cell line NCIH1568 (NSCLC, metastatic) presented higher drug response heterogeneity than RERFLCAD1 (NSCLC, primary), evidencing a more variable drug-response behaviour in the metastatic cell line. We found that 48% of cancer cell lines show >60% of drug homogeneity (SP = 0 or SP = 1) suggesting low cell variability in drug response. A total of 88% of the cell lines have a median SP > 0.7, meaning that they contain a greater number of insensitive than sensitive cells, and 17% of cell lines have a median SP > 0.9, indicating that none of the cells would exhibit sensitivity against half of the therapeutic options (Additional File 1: **Fig. S12; Table S7).** To explore the functional properties of the TCs, we correlated BCS and 12 expression programs known to be recurrently heterogeneous within cancer cell lines (RHPs). Cells enriched in the epithelial senescence-associated (EpiSen) program correlated (r > 0.6) with high sensitivity to EGFRi, in agreement with experimental validations performed by Kinker and colleagues (**Fig. 3b)**. The EMT program (EMT-II) presented higher activity in TC2 cells and correlated with high sensitivity to PI3K and HMGCR inhibitors. In contrast, TC0 and TC3 cells correlated with low EMT-II activity and high sensitivity to HDAC inhibitors, in agreement with previous studies (29, 30) (Additional File 1: **Fig. S13; Table S8**). Interestingly, EMT-high TC2 — mostly represented by brain cancer cell lines — presents a more undifferentiated transcriptional state in contrast to the rest of the therapeutic clusters (p = 8.3e−13) (30). TC2 differential expression analysis showed an up-regulated mesenchymal profile and enrichment of the EMT pathway ( Additional File 1: **Fig. S14; Table S9),** with Beyondcell results also showing a lower sensitivity to EGFRi (**Table S4**), confirming previous results where undifferentiated cell lines showed decreased sensitivity to EGFRi (**Table S10**) (31). This result shows that distinct cell functional states lead to different drug responses which are successfully detected by Beyondcell TCs. Overall, our study reveals the therapeutic landscape in multiple cancer cell lines, finding recurrent patterns of drug heterogeneity shared by specific cancer types, cell lines, as well as their relationship with functional status.

### Beyondcell characterises single-cell variability in drug response in melanoma patients

We also employed our method to study more heterogeneous samples such as primary tumour samples. For this, we used an scRNA-seq dataset from 16 melanoma patients treated with immune checkpoint inhibitors (ICIs) and 15 untreated patients. This prospective study aimed to detect transcriptional cell states related to responsiveness to ICIs and identify a transcriptional ICI-resistance program expressed by malignant cells associated with T-cell exclusion and immune evasion which predicts clinical responses to immunotherapy in melanoma patients (32). Beyondcell revealed 7 TCs, where TC2 and TC5 mainly contained cells from the untreated patients Mel79 and Mel81 respectively, whereas TC4 and TC6 included ICI-resistant patients (Additional File 1: **Fig. S15**). Non-responder patients were predicted by calculating the activation level of the ICI-resistance program with Beyondcell, thus tumour cells showing high BCS would have low response to ICIs. As expected, TC4 and TC6 defined by ICI-resistant patients showed high BCS values. TC5, which included the untreated patient Mel81, also had a high BCS so we concluded that this patient could not respond to ICIs (**Fig. 4a**). Beyondcell showed that Mel81 would respond to CDK-inhibitor (CDKi) drugs such as alvocidib, which has been proposed for use in combination with immunotherapy to improve the response in ICI-resistant patients. Conversely, ICIs in monotherapy would be the preferred treatment for Mel79 (**Fig. 4b;** Additional File 1: **Fig. S16**). This validation demonstrated that Beyondcell correctly identified non-responders to ICIs, confirming that CDKi can be a therapeutic option to overcome ICI-resistance.

**Figure 4.**
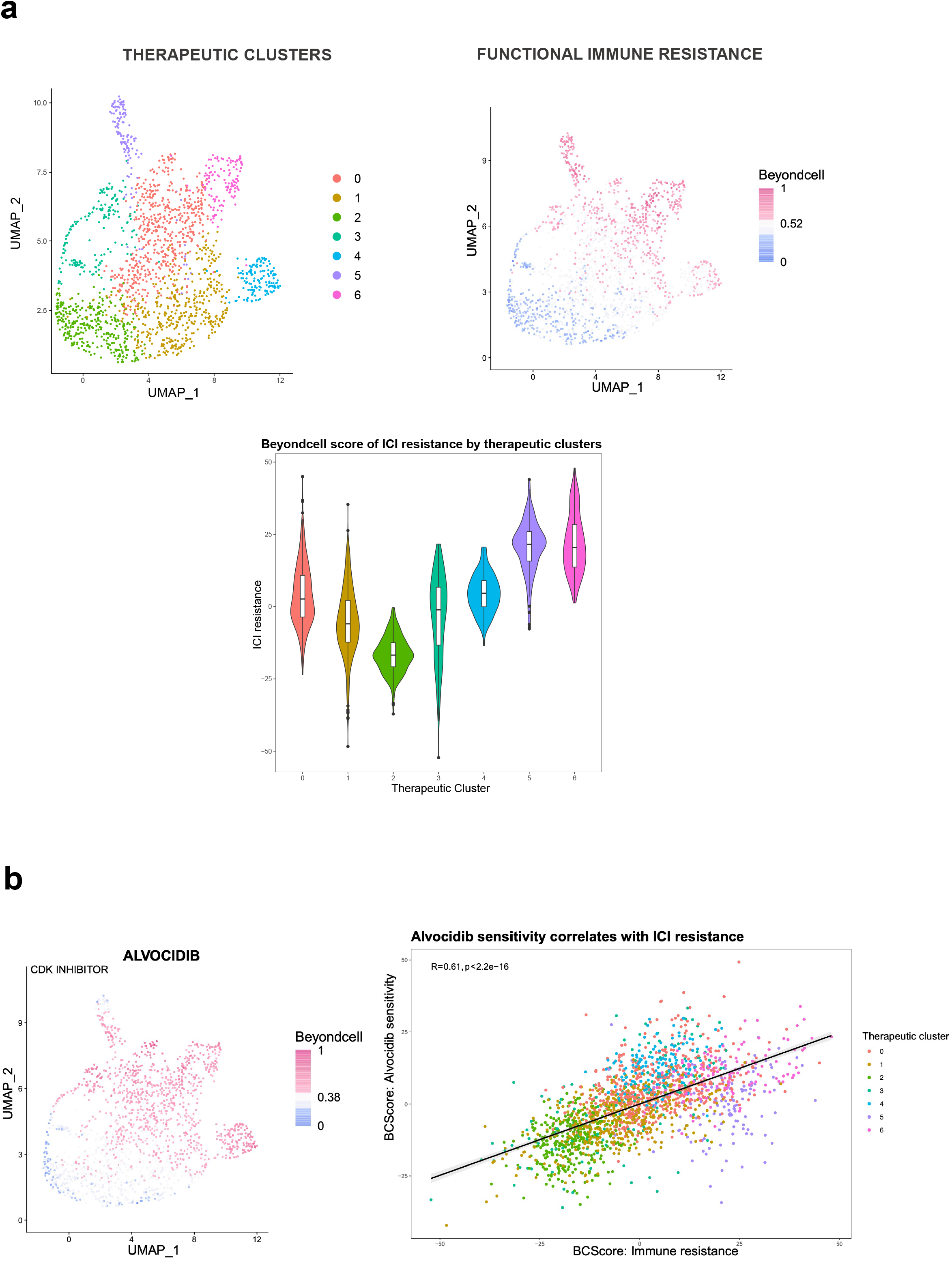
Beyondcell characterises single-cell variability in drug response in melanoma patients. **(a)** Using the Jerby-Arnon dataset, we were able to recapitulate the melanoma cells’ resistance to immune checkpoint inhibitors (ICI) at single-cell resolution. *Left:* therapeutic SS_C_ clusters, each cell is coloured according to the cluster it belongs to. *Centre:* UMAP representing Beyondcell’s predicted affinity to the functional ICI resistance signature. Higher scores indicate a higher concordance between a cell transcriptome and the signature. *Right:* A violin plot of the normalised ICI resistance score by therapeutic cluster. **(b)** Beyondcell proposes alvocidib as a therapeutic alternative to ICI cells. *Left:* UMAP of alvocidib drug score (SS_C_ collection), where higher scores indicate cells whose transcriptome is concordant with alvocidib sensitivity. *Right:* A scatterplot of alvocidib drug score and ICI resistance score. Alvocidib sensitivity positively correlates with immune resistance (R = 0.61, p < 0.01).

## Discussion

Addressing TH is a critical factor for the design of effective treatments in cancer and a current challenge for precision oncology (33). Consequently, there is a need for methodologies to detect tumour clone-specific drug sensitivities in order to properly characterise drug responses in cancer and, more importantly, to prioritise those treatments that could be clinically more effective for each patient. Here we aim to address this challenge by introducing Beyondcell, the first method to define tumour cell subpopulations of differential drug response and propose cancer-specific treatments using single-cell RNA-seq data. A key concept in Beyondcell is the “therapeutic cluster”, defined as a group of cells sharing a common drug response, which aims to address tumour therapeutic heterogeneity. For this purpose, Beyondcell uses a single-cell expression matrix and LINCS/CMap, GDSC, CCLE and CTRP drug-signature collections to calculate the Beyondcell score, an indicator for each cell of its degree of sensitivity to a drug. Then, the method also calculates the SP, a quantitative measure of the drug response homogeneity throughout the single-cell dataset reflecting the cellular variability. Finally, Beyondcell makes it possible to uncover and visualise TCs, providing a drug sensitivity ranking that prioritises the best hits to target TCs, cell subpopulations, experimental conditions or phenotypes.

As a proof of concept, we applied Beyondcell in four independent datasets comprising cancer cell lines (21,23,25) and patients (31). We observed that Beyondcell successfully identifies tumour subpopulations based on therapeutic response, and can detect differential treatments amongst a range of experimental conditions or cell clusters. In addition, Beyondcell recapitulates the biology of the datasets analysed and easily reveals sensitive, innate and acquired drug-resistant cell subpopulations in cell lines under drug exposure. Furthermore, Beyondcell is able to propose a possible treatment strategy to overcome such resistance and identifies drug-response markers (21,23). We also successfully applied Beyondcell to characterise single-cell variability in drug response in a pan-cancer cell line dataset, identifying cellular functional activities and associating them with common patterns of drug response shared by specific cancer types and cell lines (25). In addition, Beyondcell allowed us to explore tumour heterogeneity relating it to clinical drug response data to successfully predict responders and non-responders to immunotherapy in melanoma patients (32). Overall, Beyondcell allows an in-depth exploration of the TH impact in cancer therapeutics at the transcriptional level, addressing the variability of drug response in cancer cell lines and patients.

Nevertheless, the therapeutic heterogeneity revealed in our analysis pinpoints where the integration of new information is most needed. Other single-cell measurements, such as cell imaging, genetic and epigenetic state or measuring gene expression at various time points could reveal differences in the drug responses of tumour cell populations (34). Our results also reinforce the importance of tumour evolution for cancer diagnosis and treatment. Since tumour heterogeneity is related to the processes of clonal evolution, the identification of clonal populations, the clonal dynamics under selective pressure and the potential for competitive release of drug-resistant tumour subclones should be considered to prescribe more effective therapies (2). A key aspect here is to incorporate the cells’ genetic states, including information on druggable mutations and tumour vulnerabilities that could be used to target drug-tolerant cells in addition to considering the clonal evolution, in order to determine the best therapeutic opportunities to prevent or overcome tumour resistance (35).

The tumour microenvironment represents an additional layer in cancer therapeutics. Immunoediting processes — that is, the dynamic interactions between tumour cells and the immune system — drive tumour evolution and contribute to therapeutic failure (36). Several single-cell studies have suggested targeting immune cells to overcome drug resistance in some refractory tumours (37,38). Beyondcell application could be extended to the context of immune cells as a therapeutic target in cancer to study its relationship with drug treatments (39). Further short-term applications of Beyondcell would include its application to study the relationship between tumour spreading behaviour, development of metastatic phenotypes (40) and Beyondcell adaptation to the cancer single-cell spatial transcriptomics scenario (41).

## Conclusions

In summary, understanding how TH leads drug response in patients will help to design precise therapeutic regimens (single-agent, combination or sequential treatments) anticipating the appearance of relapse, managing drug resistance mechanisms, delaying tumour growth or even inducing complete tumour regression. Therefore, there is an urgent need to develop methodologies to directly address TH in the design of anticancer treatment regimens. Beyondcell provides a valuable resource for better understanding of the biological and therapeutic impact of TH. Our results highlight the utility of Beyondcell to reveal, from single-cell transcriptomics, the cellular heterogeneity in drug response in cancer cell lines and patients, identifying resistant and sensitive cellular subpopulations and proposing drugs to target them. Finally, Beyondcell has been implemented as an open-source software package that can be easily combined with scRNA-seq gold-standard methodologies in custom automated analysis pipelines. Our software is complementary to current single-cell analysis approaches, opening up a discovery space to support the design of more effective lines of therapy.

## List of abbreviations

TCs: Therapeutic clusters
TH: Tumour heterogeneity
scRNA-seq: Single-cell RNA sequencing
CCLE: Cancer Cell Line Encyclopedia
GDSC: Genomics of Drugs Sensitivity in Cancer
CTRP: Cancer Therapeutic Response Portal
LINCS: Library of Integrated Network-based Cellular Signatures
PSC: Drug perturbation signature collection
SSC: Drug sensitivity signature collection
MSigDB: Molecular Signatures Database
EMT: Epithelial-mesenchymal transition
SP: Switch Point
BCS: Beyondcell score
DSS: Drug Specificity Score
KNN: k-Nearest Neighbours
UMAP: Uniform Manifold Approximation and Projection
PCA: Principal Components Analysis
HSP: Heat Shock Proteins
RHPs: Recurrently heterogeneous programs
EpiSen: Epithelial senescence-associated
ICIs: Immune Checkpoint Inhibitors
CDKi: CDK-inhibitor

## Declarations

### Ethics approval and consent to participate

Not applicable.

### Consent for publication

Not applicable.

### Availability of data and materials

The Beyondcell algorithm is implemented in R (v. 4.0.0 or greater). All code and full documentation are available online at: https://gitlab.com/bu_cnio/beyondcell. All custom scripts used to process and analyse the data are available upon reasonable request.

The *Ben-David et al*. dataset was obtained from GEO (GSE114461). Ho *et al.*, data were downloaded from the SRA database (SRP127328). Both the pan-cancer samples from Kinker *et al*. and the Jerby-Arnon *et al*. study, were obtained from the Single Cell Data Portal website (https://singlecell.broadinstitute.org/single_cell). Samples’ re-analysis was done with the bollito pipeline (https://gitlab.com/bu_cnio/bollito).

### Author contributions

C.F-T. and M.J.J-S. designed Beyondcell, developed and maintained the Beyondcell software package. T.DD. performed the beta-testing. C.F-T., M.J.J-S., S.G-M., C.C-P. and L.G-J., executed the benchmarking analyses with input of T.DD. and supervision by G.G-L. and F.A. Analysis pipelines were implemented and optimized by C.F-T., L.G-J. and T.DD. G.G-L. and F.A. wrote the manuscript. F.A. initiated, designed, and led the study.

## Acknowledgements

We thank Dr. Thomas Walsh for reviewing the manuscript and constructive comments. We thank the whole BU staff for useful discussions. C.F-T is supported by the Paradifference Foundation. S.G.-M. was supported by Comunidad de Madrid [PEJD-2019-PRE/BMD-15732]. CNIO Bioinformatics Unit is supported by the Instituto de Salud Carlos III (ISCIII); Project RETOS RTI2018-097596-B-I00 (MCI/AEI/FEDER, UE); Spanish National Bioinformatics Institute (ELIXIR-ES, INB) Grant (PT17/0009/0011 - ISCIII-SGEFI / ERDF); Paradifference Foundation; Comunidad de Madrid (S2017/ 65 BMD-3778, LINFOMAS-CM) co-financed by European Structural and Investment Fund.

## Supplemental information

**Additional file 1:**

### Supplementary Methods

**Supplementary Figures**

Supplementary Figures 1–16.

**Supplementary Table 1**

Beyondcell results for clusters and conditions with Ho et al. dataset.

**Supplementary Table 2**

Genes differentially expressed in cluster 5 for Ho et al. dataset.

**Supplementary Table 3**

Cell distribution in clusters, cancer types and cell lines for Kinker et al. dataset.

**Supplementary Table 4**

Differential sensitivity drugs in cancer type and clusters for Kinker et al. dataset.

**Supplementary Table 5**

Beyondcell drug scores correlation modules for Kinker et al. dataset.

**Supplementary Table 6**

Differential EGFRi vulnerability analysis for cell lines in TC1 in comparison to TC2.

**Supplementary Table 7**

Cell lines drug homogeneity for Kinker et al. dataset.

**Supplementary Table 8**

Correlation of drug sensitivity and Recurrent Heterogeneous Programs (RHP) Beyondcell scores.

**Supplementary Table 9**

Differential gene expression analysis of TC2 cells vs rest of TCs.

**Supplementary Table 10**

Differential EGFRi vulnerability analysis for undifferentiated cell lines in comparison to differentiated cells.

