## SupplementaryFigures_and_Methods for "Beyondcell: targeting cancer therapeutic heterogeneity in single-cell RNA-seq"

### Additional File 1

#### Supplementary Figures

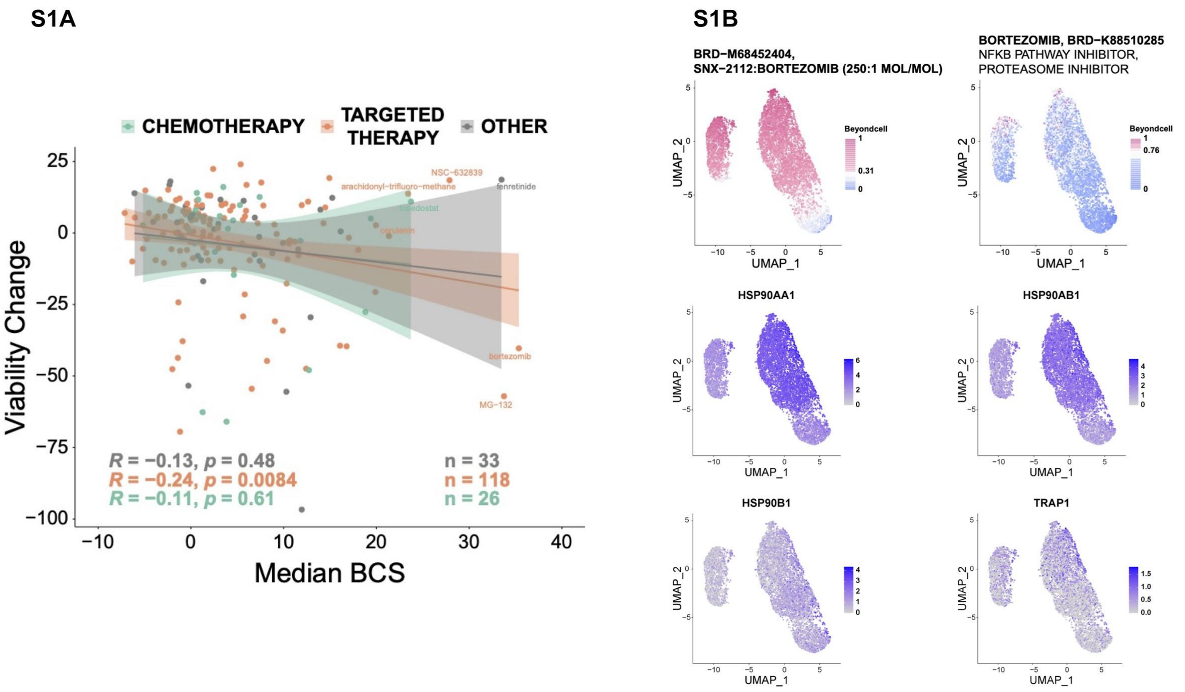


**Supplementary Figure 1. Beyondcell score correlation with cell viability and prediction of response using a bortezomib signature. (A)** Median BCS obtained with $PS_C$ signatures for cells at t0 correlate negatively with the viability measures reported by Ben-David *et al*. BCS values were regressed to remove the effect of the number of genes detected in each cell and, to avoid duplicated measurements, we computed the mean BCS and mean viability score for each compound. Drugs were then classified into three groups: chemotherapy, targeted therapy and others (including immunotherapy, hormone therapy and photodynamic therapy). Only the BCS for targeted therapies showed a significant correlation with MCF7-AA cell viability (R = -0.24, p = 8.4e-03). **(B)** Predicted sensitivity of MCF7-AA cells in response to bortezomib and bortezomib combination signatures ($SS_C$). Cells are predicted to be insensitive to bortezomib alone (sig_21310) but sensitive to bortezomib in combination with SNX-2112 (sig_21560). The longer the time for which MCF7-AA cells have been exposed to bortezomib, the more insensitive to sig_21560. Sensitivity to bortezomib:SNX-2112 could be explained by the higher expression of cytosolic Heat Shock Proteins (*HSP90AA1* and *HSP90AB1*), which are targets of SNX-2112. Other non-cytosolic HSP are shown for comparison. *HSP90AA2* was not present in the original Seurat object.

**S2**

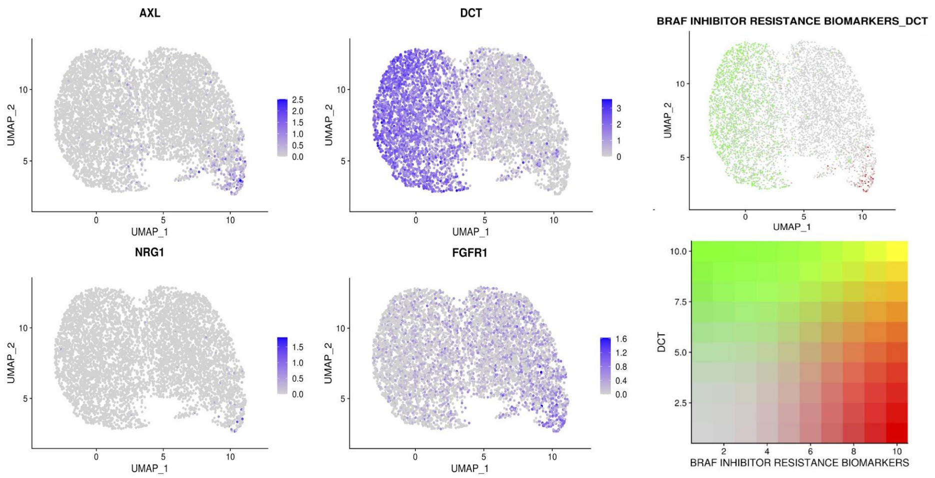


**Supplementary Figure 2. Analysis of resistance biomarkers in the melanoma cells from Ho *et al*. dataset.** *Left:* UMAP plots of Beyondcell object obtained using $SS_C$ drugs signatures coloured according to the expression of some BRAF inhibitor-resistant biomarkers: *AXL*, *NRG1*, *DCT* and *FGFR1*. *Right*: UMAPs showing the coexpression of both *DCT* and the sum expression of BRAF inhibitor-resistance biomarkers (*JUN, WNT5A, PDGFRB, EGFR, NRG1, FGFR1* and *AXL*).

**S3**

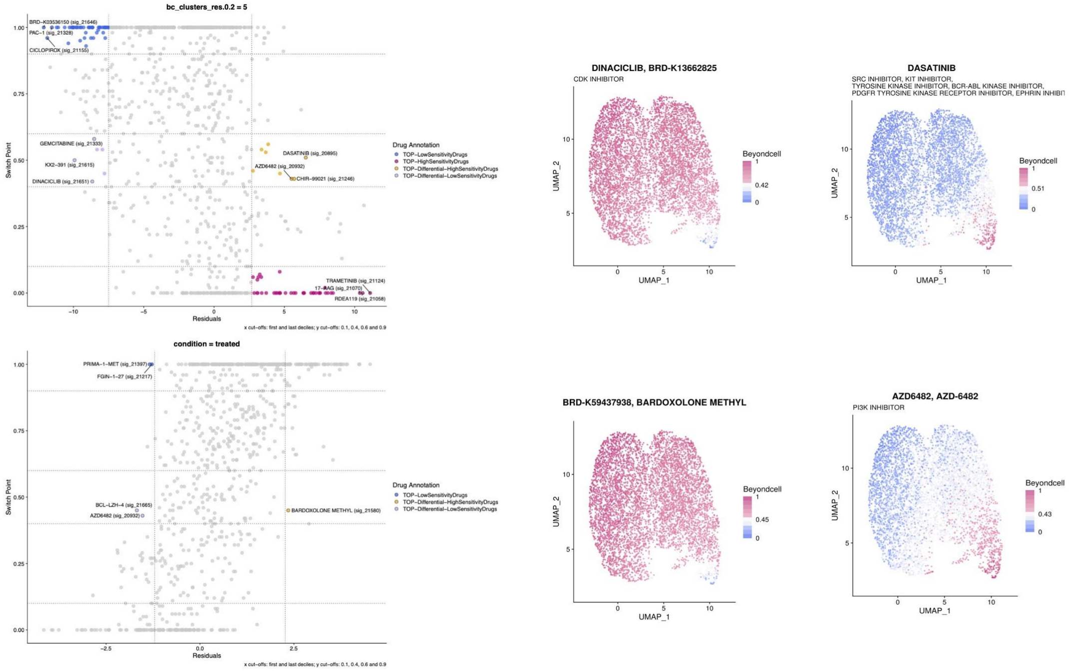


**Supplementary Figure 3. Beyondcell selects specific drugs for a cluster or group of cells.** *Left* 4 squares plots of distinct cell groups and UMAPs showing the BCS of drugs to which the selected cells show differential sensitivity. *Right Top*: cells in TC5 are differentially sensitive to dasatinib (SRCi) and unresponsive to dinaciclib (CDKi) and gemcitabine. *Right Bottom*: cells in resistant condition are differentially sensitive to bardoxolone methyl (NF-kBi) and unresponsive to AZD6482 (PIK3i).

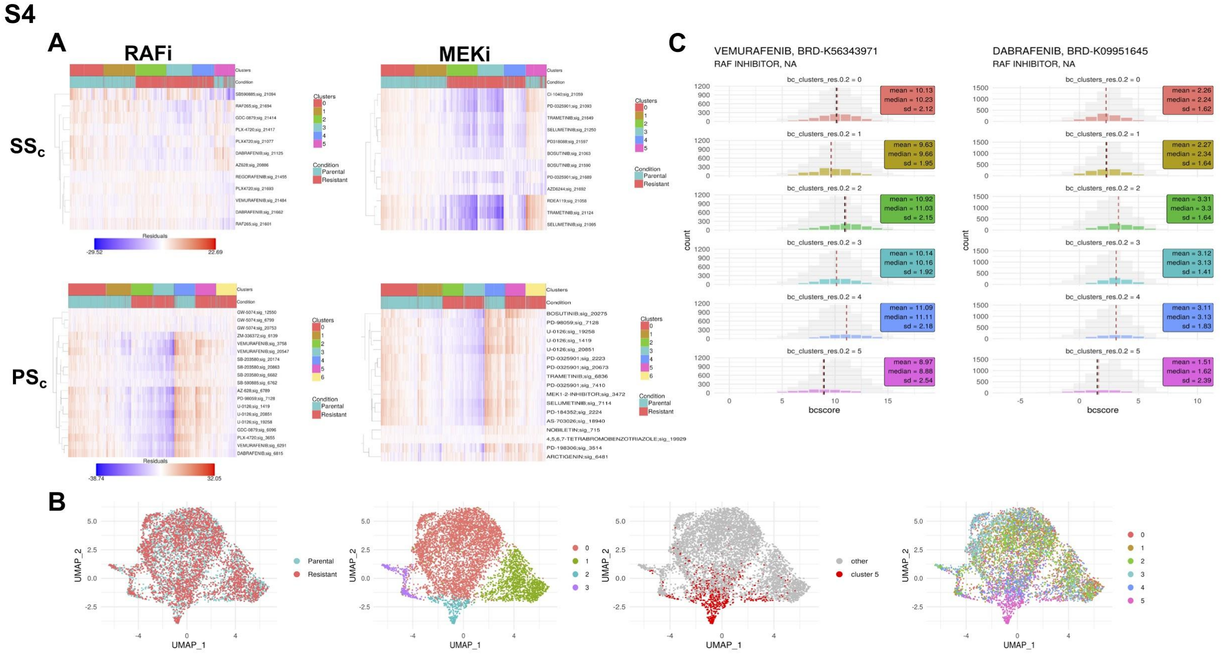

**Supplementary Figure 4. Beyondcell classifies cells in therapeutic clusters with different BRAF inhibitor responses. (A)** Heatmaps showing the Beyondcell score residuals of *BRAF* and MEK inhibitors from $PS_C$ and $SS_C$ signatures in each single cell. Heatmaps are ordered by each therapeutic cluster and the treatment condition of each cell is annotated. **(B)** UMAP plots based on the expression data. *From left to right:* treatment condition, Seurat clusters, the therapeutic cluster 5 and Beyondcell's therapeutic clusters. **(C)** Histogram representing vemurafenib and dabrafenib Beyondcell scores in each therapeutic cluster computed with $SS_C$ drug signatures. TC5 shows lower sensitivity to RAF inhibitors than the rest of the clusters.

**S5**

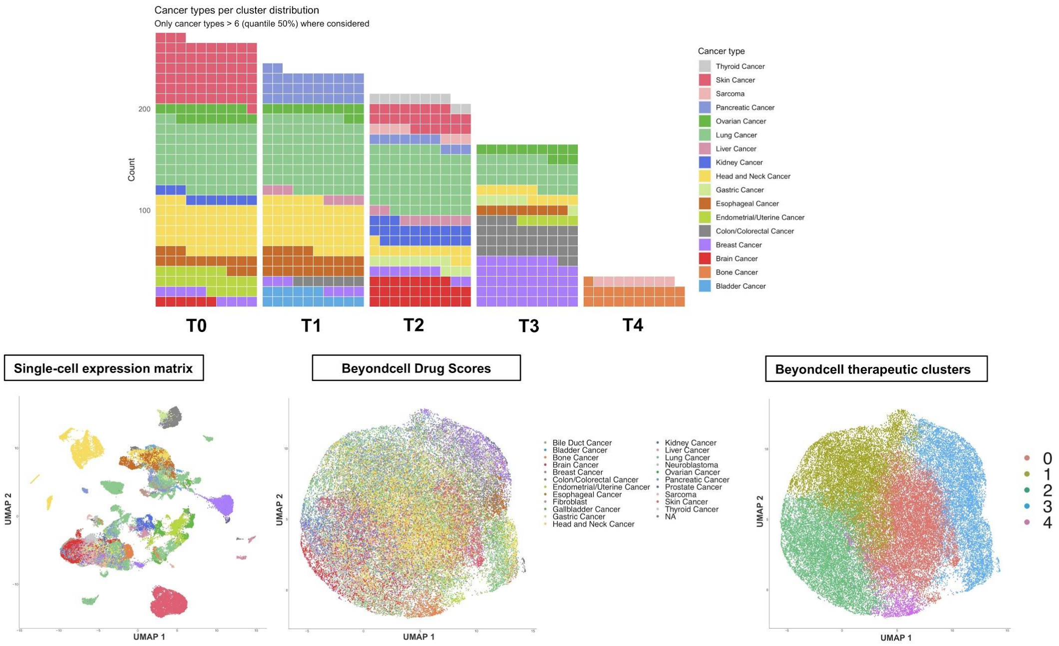


**Supplementary Figure 5. Therapeutic landscape in pan-cancer cell line data.** *Top:* We have found 5 therapeutic clusters that sum up all the possible drug responses in this cohort. The waffle plot shows the cancer type proportions of each cluster. *Bottom:* The UMAP shows the differences in the heterogeneity based on whether we are analysing the expression or the drug susceptibility profiles.

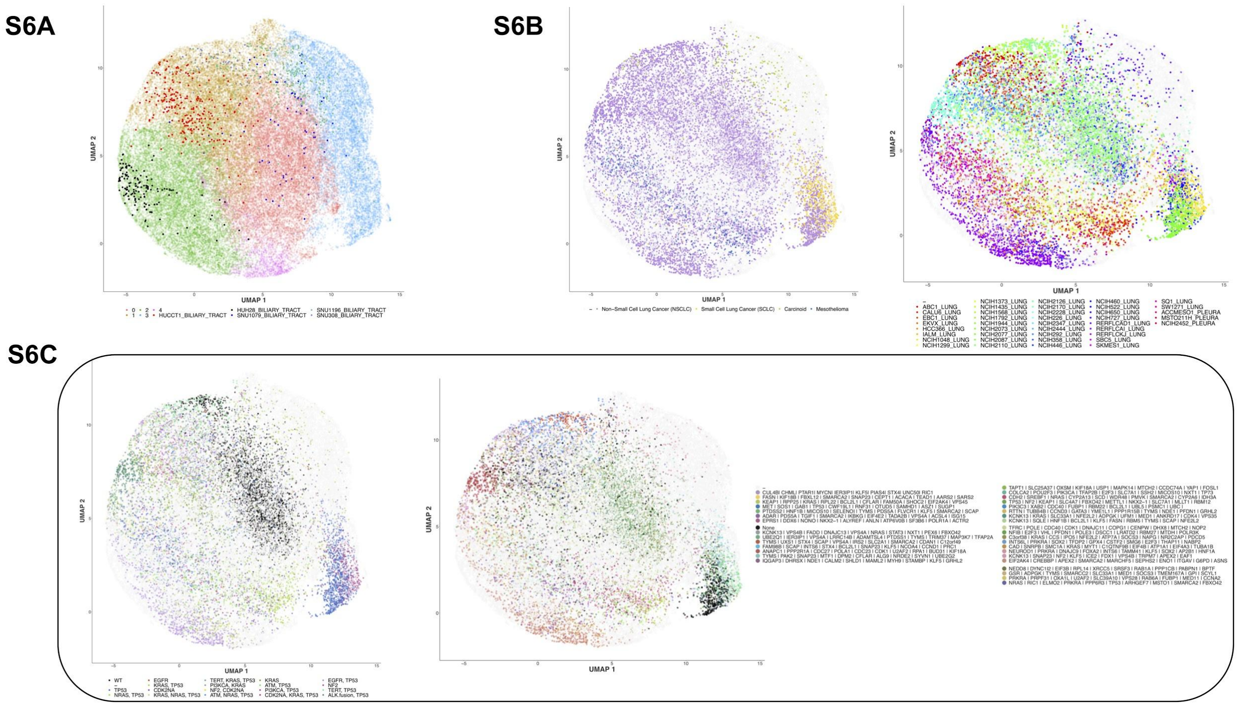


**Supplementary Figure 6. (A)** UMAP highlighting bile duct cancer cell lines. HUCCT1, HUH28, SNU1079 and SNU1196 are clustered in different TCs (TC0, TC1, TC2 and TC3 respectively) showing distinct drug response profiles despite belonging to the same cancer type. **(B)** The UMAP of the lung cancer cell lines shows the diversity in their drug responses as they  are distributed between the TCs regardless of their classification in squamous or adenocarcinoma subtypes. **(B)** Lung cancer cell lines are grouped by drug response and not driven by other dependencies. *Left:* The UMAP shows lung cancer cell lines TCs coloured by their oncogenic drivers. *Right:* The UMAP depicts the gene dependencies in lung cancer cell lines.

**S7**

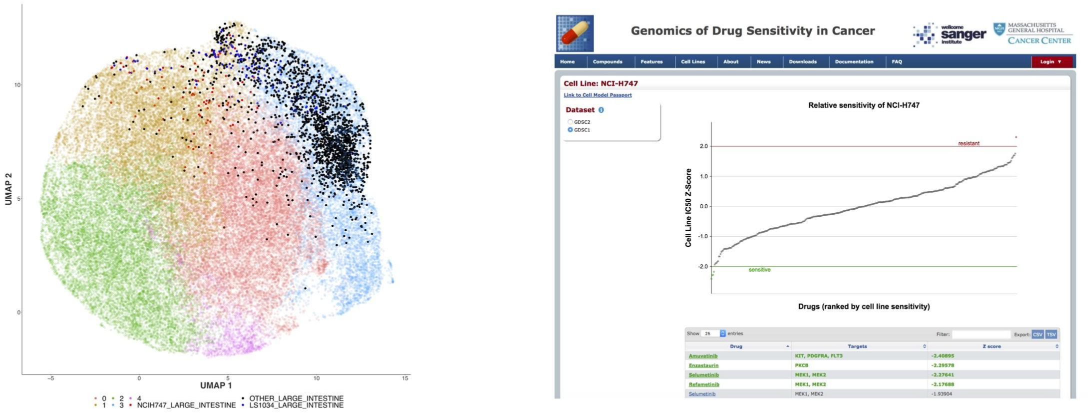


**Supplementary Figure 7**. **UMAP highlighting NCIH747 cancer cell line.** NCIH747 colon/colorectal cancer cell line located in TC1 with an experimental sensitive response to selumetinib (MEKi) from GDSC portal.

**S8**

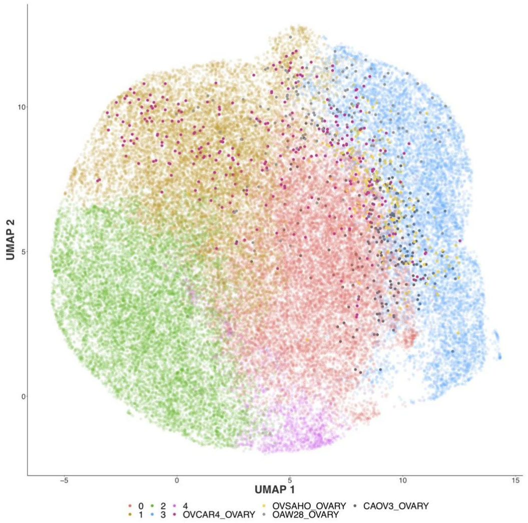


**Supplementary Figure 8**. **UMAP highlighting ovarian cancer cell lines.** The UMAP shows cells belonging to five high-grade serous ovarian cancer cell lines.

S9

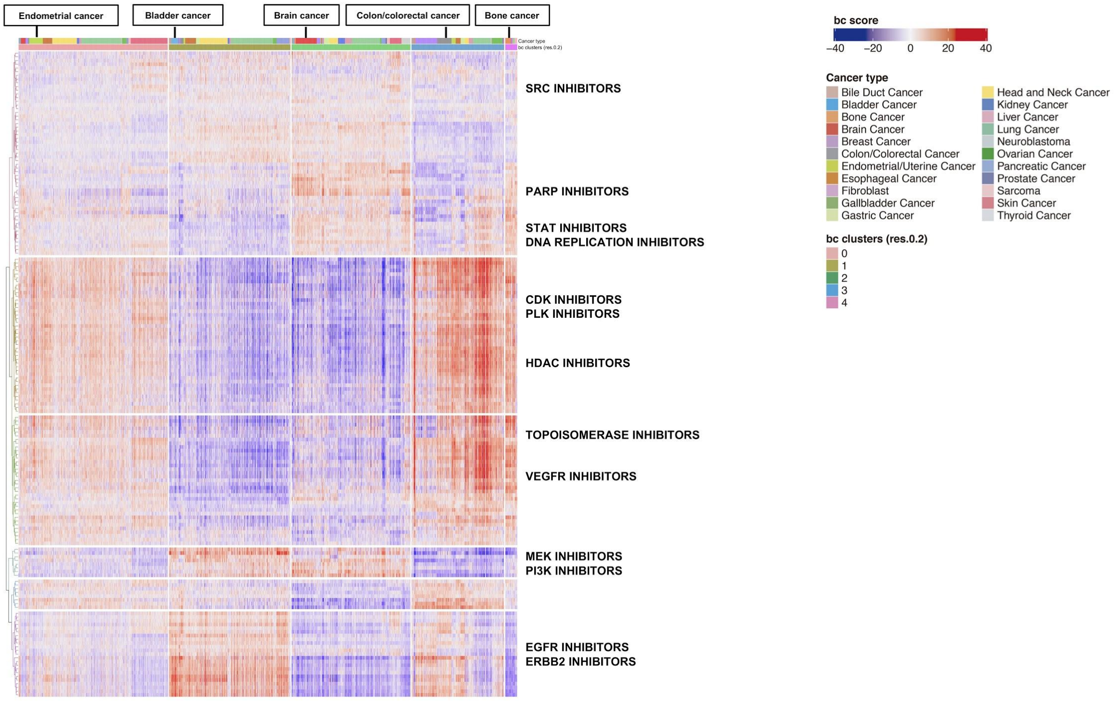


**Supplementary Figure 9**. The heatmap shows both the cluster and cancer-specific drugs with a heterogeneous sensitivity pattern. A total of 130 targeted therapies have been considered.

S10

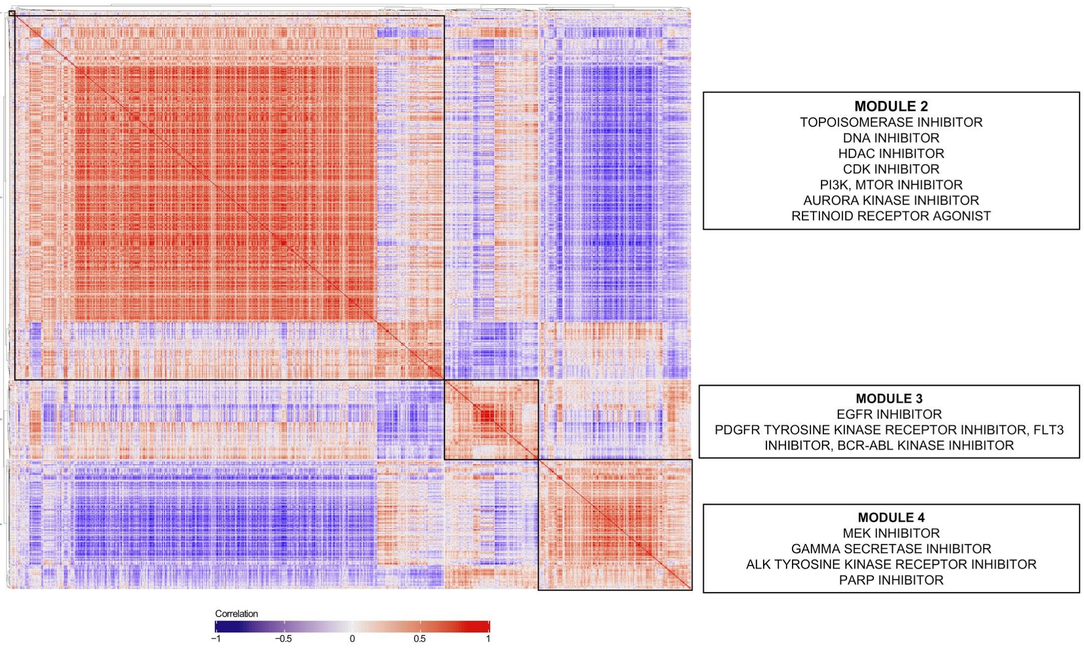


**Supplementary Figure 10**. The heatmap depicts all pairwise correlations between the BCS, ordered by hierarchical clustering. Three main clusters were found, associated with distinct MoA patterns. Further information can be found in Supplementary Table 5.

## S11

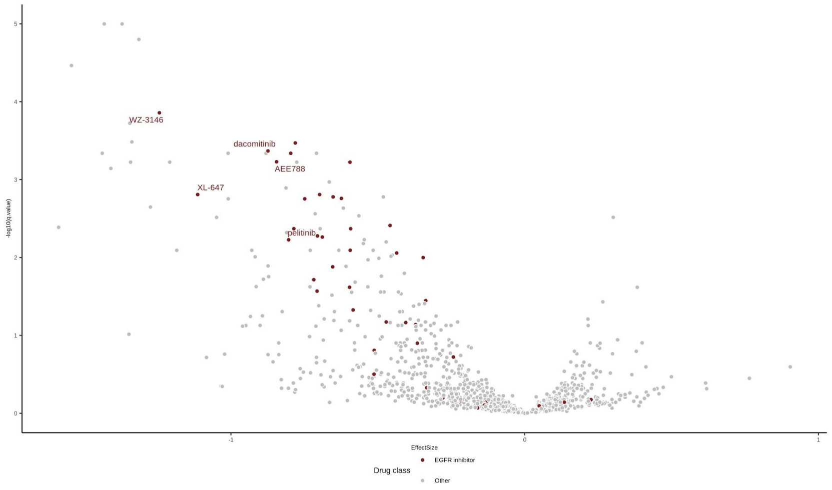


**Supplementary Figure 11**. Differential drug vulnerability analysis showed increased sensitivity to EGFR inhibitors for cell lines in TC1 in comparison to TC2 which shows decreased sensitivity. A volcano plot of the differential drug effect size and the adjusted p-value for each drug comparison. EGFR inhibitors are highlighted in red against the rest of the compounds found in the PRISM dataset.

## S12

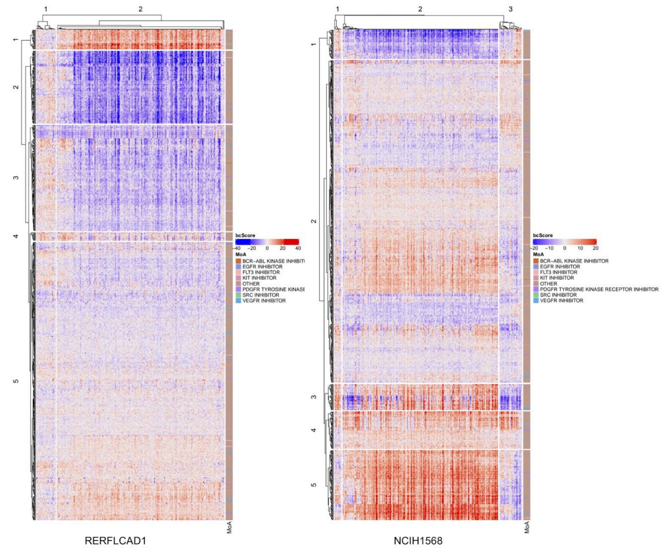


| Cell line | Homogeneous sensitivity | Homogeneous resistance | Total heterogeneity |
|---|---|---|---|
| *RERFLCAD1* | 20.66% | 47.24% | 66.1% |
| *NCIH1578* | 17.03% | 38.25% | 58.28 |

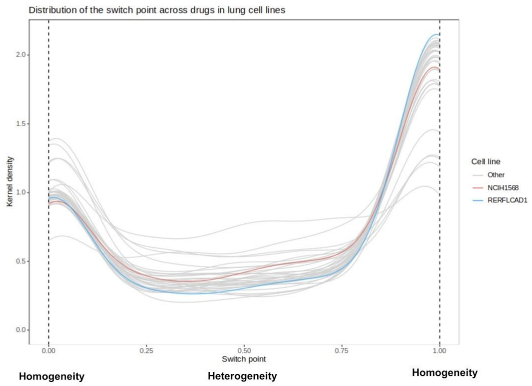


**Supplementary Figure 12**. Drug response homogeneity in *RERFLCAD1* (primary malignancy) and *NCIH1578* (metastatic) lung cancer cell lines at single-cell resolution. *Left:* Heatmaps of the normalised drug scores for the $SS_C$ drug collection. Individual cells (columns) were grouped by consensus clustering after 1000 iterations. Drugs (rows) were hierarchically clustered on the Euclidean distance between drug scores. *Right:* Distribution of the switch point in lung cancer cell lines across the $SS_C$ drug collection. RERFLCAD1 and NCIH1568 are highlighted in light blue and red, respectively.

**S13**

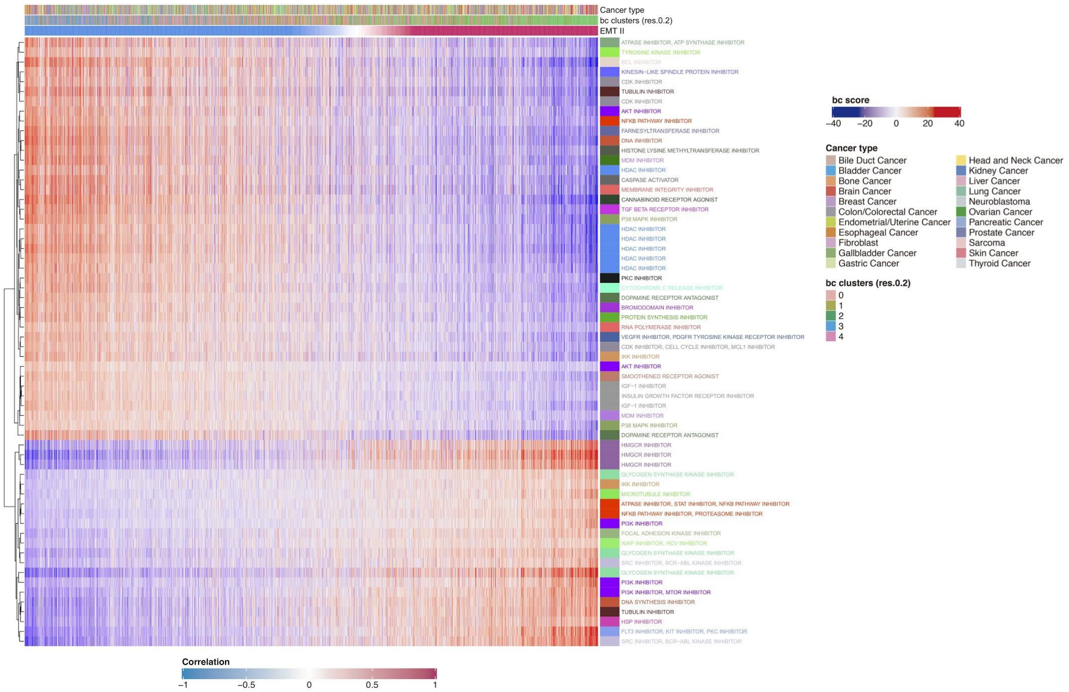


**Supplementary Figure 13**. We selected drugs with an absolute correlation > 0.6 with the EMT II program. A hierarchical clustering was applied to the rows. Columns were ordered by their correlation degree. HDAC inhibitors mainly correlated with the EMT II *low*, while PI3K inhibitors or *HMGCR* had a higher correlation with cells expressing EMT II.

**S14**

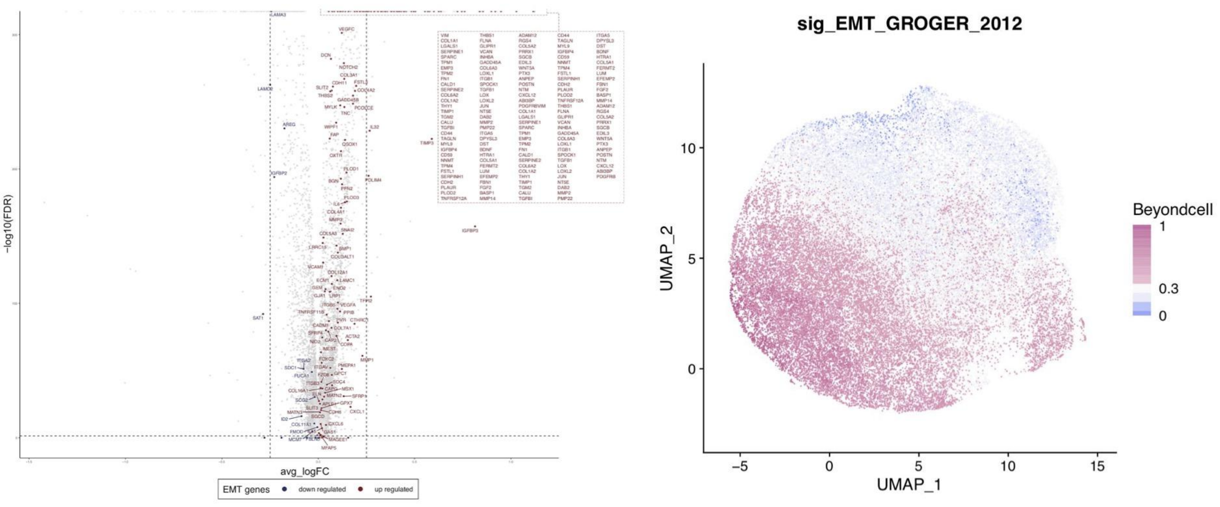


**Supplementary Figure 14**. TC2 differential expression analysis showed an up-regulated mesenchymal profile and enrichment of the EMT pathway. *Left:* A volcano plot of the average log fold change and the adjusted p-value for each gene. EMT related genes have been annotated depending on whether they are upregulated (red) or downregulated (blue). Right: A UMAP plot showing the BCS for the EMT program.

**S15**

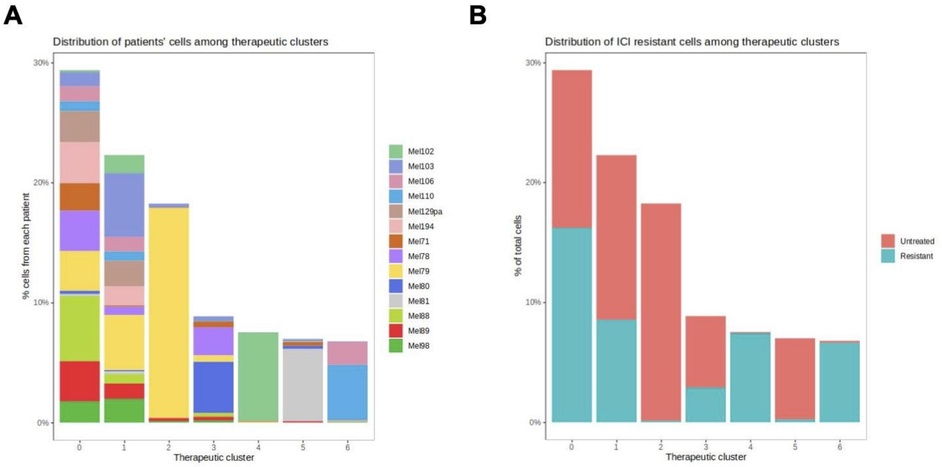


**Supplementary Figure 15. Distribution of malignant cells across therapeutic clusters from Jerby-Arnon *et al*.** **(A)** A bar plot illustrating the proportion of malignant cells from each melanoma patient. **(B)** An identical bar plot showing the relative composition of untreated and ICI-resistant cells for each cluster.

**S16**

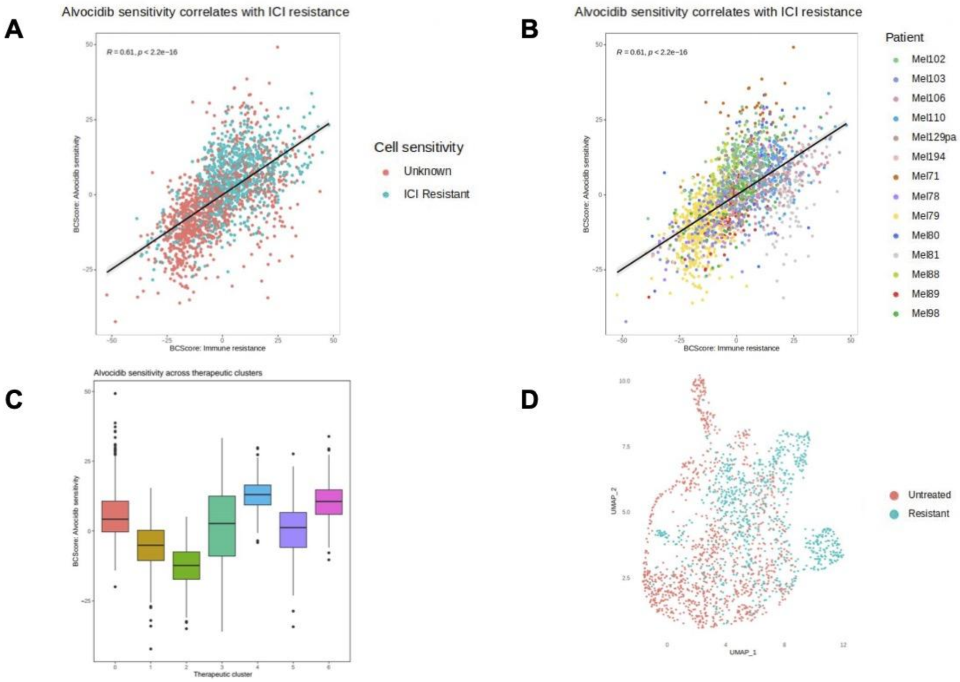


**Supplementary Figure 16. Alvocidib sensitivity correlates with ICI resistance**. **(A)** The scatter plot of figure 2D with ICI-resistant cells in light blue. **(B)** The same scatter plot, labeled by patient of origin. **(C)** A box plot of Beyondcell's alvocidib drug score by therapeutic cluster. **(D)** The same UMAP from figure 2C with ICI-resistant cells in light blue.

### Supplementary Methods

#### Data description

The Library of Integrated Network-based Cellular Signatures (LINCS, http://www.lincsproject.org) (Subramanian et al., 2017) is a catalogue of gene expression data associated with cell lines exposed to a variety of perturbing agents, such as small molecules (about 5500), FDA drugs (~1300) and shRNA silencing (22,119 genetic perturbagens). LINCS, based on L1000 high throughput technology, is an extension of the Connectivity Map, which has been successfully used for drug repositioning (Lamb et al., 2006). From the ~20K small molecules tested in the LINCS L1000 dataset, only those with an identifiable common drug name were selected. This reduced the number of drugs to 4690. In the CCLE (Barretina et al., 2012) 28 drugs were tested and the mRNA expression of 1037 cell lines were profiled using Affymetrix U133Plus2 arrays. The gene-centric RMA-normalised mRNA expression data, the cell line information and the drug response data from Cancer Cell Line Encyclopedia were downloaded from the CCLE data portal (https://portals.broadinstitute.org/ccle/home). In the CTRP (Basu et al., 2013), 545 drugs (single or combination) were tested against 887 cell lines. We downloaded the drug response data of the CRTP 2.1 project from the CTD2 Data Portal (https://ocg.cancer.gov/programs/ctd2/data-portal). The gene expression data used in order to obtain the CTRP expression signatures were those of the CCLE (see above). Of the 887 cell lines, 805 were present in the CCLE dataset. In the GDSC data (Garnett et al., 2012), 265 compounds corresponding to 250 different drugs were tested against 1074 cell lines. Cell lines were profiled for gene expression using the Affymetrix Human Genome U219 array. Expression data were normalised using RMA (Irizarry et al., 2003). Drug response data and preprocessed mRNA gene expression data were downloaded from the GDSC web portal (http://www.cancerrxgene.org/downloads).

#### Signature generation

A gene expression signature is a general model for the representation of the transcriptional changes associated to a given biological phenotype or perturbation. In this study we have considered two expression signature collections: the $PS_C$ captures the transcriptional changes induced by a drug; while the $SS_C$ captures the sensitivity to the drug effect. In both cases, these expression signatures are obtained from a differential expression analysis, although using different designs. Despite its different biological interpretations, both types of signature were represented as the two gene sets formed by the *N* most up and down-regulated genes (the *UP* and *DN* genesets). Several *N* were tested (50, 100, 250 and 500) and no significant disagreement was found between the top synergistic and antagonistic interactions (data not shown). Thus, following (Iorio et al., 2010), *N* was taken as 250.

##### Drug perturbation signature collection ($PS_C$)

Drug-induced expression signatures were obtained from experiments in which the transcriptional state of the cell is measured before and after treatment with the drug. This makes it possible to study the transcriptional effect of the drug. In order to obtain consensus expression signatures for each drug, a differential expression analysis was performed on control vs treated cells using *limma* (Ritchie et al., 2015). In the LINCS L1000 data, all the wells in which the same drug was used were considered as treated samples. All the DMSO-treated wells from all the plates with at least one treated well were considered as untreated controls. The plate in which the drug was tested was taken as a covariate in the expression analysis. As only a type of cell line is used in each plate, using this covariate we take into account both the technical batch, due to different plates, and the biological variability, due to different cell lines. Different drug concentrations and exposure time to the drug were not taken into account. In the LINCS L1000 data some drugs (*pert_iname*) are represented by different molecules, (pert_id) usually from different vendors. In these cases, we obtained the pert_id associated signatures, that is, associated to the molecule, and a consensus signature in which all the pert_id corresponding to the same drug were considered. In this last case, the pert_id was also taken as a confounding variable. The *t-moderate*d statistic was used as a measure of the expression of the gene. It was preferred over the logFC because the *t-statistic* takes into account the sampling variance. However, both statistics were highly correlated in all the signatures tested.

##### Drug sensitivity signature collection ($SS_C$)

Drug-induced expression signatures were obtained from pharmacogenomics experiments in which the pharmacological response to a drug and the transcriptional state before treatment were considered. In order to obtain the expression signature, a differential expression analysis against a measure of drug response was performed using *limma.* The area under the curve (AUC) was used as the measure of drug response, as contrary to $IC_{50}$, it can always be estimated without extrapolation from the dose-response curve, and also because it has shown more accuracy in the prediction of drug response (Jang et al., 2014).

In GDSC 2.0, the AUC was taken as provided in the original dataset. In order to make the AUC from the CTRP dataset comparable to that of the GDSC, the AUC was recalculated from raw data, normalised and then truncated at 1 in order to keep the values in the interval (0,1). AUC calculation was carried out using the *drc* R package (Ritz et al., 2015). In the CCLE dataset, area under the curve was calculated from the related Activity Area considered in the data according to:

$$AUC = \frac{10 - ActArea}{10} \quad (1)$$

As the AUC is inversely related to drug response, it was taken with negative sign for the differential expression analysis.

In all the projects considered, the tumour origin of the cell lines were considered as confounding variables. In the CCLE the covariate was created by combining the *Site.Primary* and the *Histology* information of the cell line. In the GDSC 2.0, the variable *Site* and in the CTRP, the '*CCLE primary Site*' was used.

##### Beyondcell score calculation

The Beyondcell score evaluates the activity of a signature of interest in a single-cell RNA-seq experiment. The transcriptomics data needs to be pre-processed, meaning that proper cell-based quality control filters, as well as normalisation, scaling and clustering of the data, should be applied prior to the analysis with Beyondcell. When analysing a bidirectional gene signature (a signature with separate sets of up-regulated and down-regulated genes), the Beyondcell score is independently obtained for each signature mode. The individual sum of the expression is calculated and divided by the number of genes in the given signature that are present in the single-cell expression matrix. The obtained raw scores are normalised and the individual up and down normalised scores are summed and rescaled between 0 and 1. In cases where the gene signature is unidirectional, all steps remain the same, although the rescaling will only be applied to one set of normalised scores.

More formally:

Let $X = (x_{ij})$ be a single-cell expression matrix with $n$ genes contained in $I = \{i_1, \dots, i_n\}$ and $m$ cells contained in $J = \{j_i, \dots, j_m\}$. Also, consider we have a geneset $GS_M = \{S_{M,1}, \dots, S_{M,p}\}$ which consists in a set of *p* sets of genes (signatures, denoted as $S$ ) per mode in $M = \{UP \vee DN\}$.

To compute the BCS, first we calculate the intersection between $I$ and each $S_M$ in $GS_M$:

$$G_{M,S_M} = I \cap S_M \quad for\ S_M\ in\ GS_M \quad (2)$$

Then, we obtain the submatrix of $X$ that contains only the genes present in $G_{M,S_M}$. We denote this submatrix as $Y = (y_{gj})$, with $q$ genes contained in $G_{M,S_M} = \{g_1, \dots, g_q\}$. The raw score for each cell $j$ and signature $S_M$ within a mode $M$ is equal to the mean expression of the genes belonging to $G_{M,S_M}$.

$$raw_{M,S_M,j} = \bar{y_j} = \frac{1}{|G_{M,S_M}|} \cdot \sum_{k=1}^{q} y_{kj} \quad (3)$$

Furthermore, we normalise the raw scores using the mean and the standard deviation of the gene expression.

$$norm_{M,S_M,j} = raw_{M,S_M,j} \cdot \frac{\sum_{k=1}^{q} y_{kj} - \sqrt{\frac{\sum_{k=1}^{q}(y_{kj}-\bar{y_j})^2}{q-1}}}{\bar{y_j} + \sqrt{\frac{\sum_{k=1}^{q}(y_{kj}-\bar{y_j})^2}{q-1}}} \quad (4)$$

Finally, we operate on the normalised scores and scale the results between $[0,\ 1]$ for each signature $S_M$.

$$BCS_{S_M,j} = \begin{cases} (norm\ _{UP,S_M,j} - norm\ _{DN,S_M,j})\ [0,1] & \text{if } M = \{UP, DN\} \\ \ \ norm\ _{UP,S_M,j}\ [0,1] & \text{if } M = \{UP\} \\ -norm\ _{DN,S_M,j}\ [0,1] & \text{if } M = \{DN\} \end{cases} \quad (5)$$

In general, the Beyondcell score aims to evaluate the susceptibility of each cell based on the analysed gene signature. A high score can be interpreted as the concordance between the signature and the analysed cell, while a low score can be interpreted as a discordance between them.

During the rescaling process, a *switch point* per signature ($SP_{S_M}$) is calculated in order to determine the value where negative scores *switch* to positive scores and *vice versa*. When analysing unidirectional signatures, the SP will always be 0 in up-regulation signatures ($M = \{UP\}$) and 1 in the case of down-regulation signatures ($M = \{DN\}$). For bidirectional signatures ($M = \{UP,\ DN\}$), the SP is calculated as follows:

$$SP_{S_M} = \begin{cases} BCS_{S_M,r}\ [0,1] & \text{if } \exists r | BCS_{S_M,r} = 0 \\ \left(\frac{\max BCS^{-}_{S_M,j}\ +\ \min BCS^{+}_{S_M,j}}{2}\right)\ [0,1] & \text{if } \nexists r | BCS_{S_M,r} = 0 \end{cases} \quad (6)$$

Being $BCS^{-}_{S_M,j} = max(BCS_{S_{M,j}},\ 0)$ and $BCS^{+}_{S_M,j} = max(-\ BCS_{S_{M,j}},\ 0)$ the positive and negative parts of $BCS_{S_M,j}$.

- If there is a normalised BCS that is equal to 0 after summing up and down non-regressed normalised scores ($BCS_{S_M,r} = 0$), the $SP_{S_M}$ is equal to the corresponding scaled score (regressed or non-regressed).
- If there isn't any normalised BCS that is exactly 0, the $SP_{S_M}$ is computed using the value closer to 0 (arithmetic mean of the maximum negative normalised score and the minimum positive normalised score).

When visualizing the scaled scores, the SP therefore, will allow us to detect cell-based changes in the regulatory processes concerning the signatures of interest. As an example, an all-sensitive dataset will have SP equal to zero, as there won't be any negative scores for that specific drug in the whole population. Conversely, a dataset insensitive to a certain drug, will be expected to have an SP close to 1. Intermediate SP, as a contrast, will indicate that the dataset contains both susceptible and non-susceptible cells.

#### Expression threshold

Some $G_{M,S_M}$ might contain a high number of genes $g$ with 0 expression ($y_{gj} = 0$). Due to limitations in single-cell technologies, currently we are not able to distinguish between a biological zero ($g$ has truly no expression) or a technical zero (uncaptured mRNA).

In order to correct for this, we have introduced an expression threshold $t$ in the computation of BCS. $t$ is the minimum proportion of genes in $G_{M,S_M}$ that must fulfil $y_{gj} > 0$. If the proportion of genes with non-zero expression is below $t$, the raw and normalised scores are reported as $NaN$.

$$raw_{M,S_M,j} = norm_{M,S_M,j} = NaN \quad if\ |\{g \in G_{M,S_M} | y_{gj} > 0\}| < t \cdot 100 \tag{7}$$

Thus, if any $norm_{M,S_M,j} = NaN$, the final BCS for that cell and signature will be:

$$BCS_{S_M,j} = \begin{cases} norm_{UP,S_M,j}\ [0,1] & \text{if } M = \{UP, DN\} \wedge norm_{UP,S_M,j} \neq NaN \wedge norm_{DN,S_M,j} = NaN \\ -norm_{DN,S_M,j}\ [0,1] & \text{if } M = \{UP, DN\} \wedge norm_{UP,S_M,j} = NaN \wedge norm_{DN,S_M,j} \neq NaN \\ NaN & \text{if } \forall norm_{M,S_M,j} = NaN \end{cases} \tag{8}$$

#### Data sets processing

In the dataset from *Ben-David et al*., an already processed matrix was obtained from GEO (GSE114461). MCF7-AA cells expressing between 1500 and 6000 genes were kept. Data were normalised using *Seurat*'s standard normalisation (Satija et al., 2015) and cell cycle effect was regressed out. The first 25 components were selected for the downstream clustering analysis. We integrated the data, but no distinct treatment-based cluster was found (**Fig. 2a, right**).

For Ho *et al*., data were downloaded from the SRA database (SRP127328). A reanalysis of the samples was applied using the bollito pipeline (https://gitlab.com/bu_cnio/bollito). Cells expressing between 1800 and 9500 genes were kept. Samples were normalised using *Seurat*'s standard normalisation and a linear dimensionality reduction was performed using Principal Components Analysis (PCA) on the expression data. We used 12 principal components to cluster the cells and visualised the identified clusters by using a UMAP.

In the case of the pan-cancer samples from Kinker *et al*., we downloaded the UMI counts table, obtained from the Single Cell Data Portal website (https://singlecell.broadinstitute.org/single_cell) and processed it using *Seurat* (v. 3.2). Only those cells with a number of detected genes ranging from 2000 to 9000 were retained, as described in the original paper. Samples were normalised using *Seurat*'s standard normalisation and a linear dimensionality reduction was performed using PCA on the expression data. We used 10 principal components to cluster the cells and visualised the identified clusters by using a UMAP.

Finally, for the Jerby-Arnon *et al*. study, the UMI counts table was also obtained from the Single Cell Data Portal website. Only malignant cells were retained for the expression analysis. Cells expressing between 2000 and 6000 genes were kept. Cells were normalised and a clustering analysis was applied using the first 20 components.

#### Beyondcell analysis

For each dataset, we obtained the BCS for both the $SS_C$ and $PS_C$ as well as a selection of pathways. For each drug signature, only the top 250 genes were considered. The expression threshold to compute BSC was set at a minimum of 10% of the genes comprising each signature. After computing the BCS, we regressed the specific unwanted sources of variation belonging to each dataset using the *bcRegressOut* function. Then, a clustering analysis was applied to the regressed scores. Once the clusters were identified, we computed the BCS statistics for each therapeutic cluster and each treatment group with the *bcRanks* function. Both the BCS and the gene expression matrices were visualised using a UMAP. Therapeutic clusters were identified by using the *bcClusters* function. The *bcSignatures* function was used in order to visualise the selected drug signatures as well as expression of biomarkers. Moreover, we have applied the function *bcHistogram* to assess the differences in sensitivity between therapeutic clusters for a selection of signatures and the function *bc4Squares* to provide a visual ranking of the signatures according to the residuals and SP. All these functions are implemented in the Beyondcell package.

In the case of Ben-David *et al*. data, we regressed the number of detected genes in MC7-AA cells. Then, we computed a UMAP projection with the first 10 principal components at a resolution of 0.1. Using *bcClusters* we identified 4 TCs that overlapped with the time-points (t0, t12, t48 and t96) after exposure to bortezomib (**Fig. 2b, centre and right**). Moreover, we used *bcSignatures* to visualise the *Beyondcell* scores of bortezomib signatures such as sig_18868 (**Fig. 2b**) or sig_21560 and bortezomib combinations like sig_21310 (**Fig. S1B, top row**). In addition, we interrogated the expression of genes encoding for Heat Shock Proteins (HSP) using this same function (**Fig. S1B, centre and bottom rows**). Finally, we computed the Pearson correlation between the median BCS of cells at t0 and the viability change of MCF7-AA cells reported by Ben-David *et al*. In **Fig. 2a** we plotted the non-regressed Beyondcell scores obtained using $SS_C$ signatures, whereas in **Fig. S1A** we represented the regressed BCS calculated from $PS_C$ signatures. To obtain the latter figure, we first computed the mean viability change per drug and the mean regressed BCS score per drug and cell. We did so in order to remove duplicated points for the same compound. Then, drugs were classified into three groups: chemotherapy, targeted therapy and others (including immunotherapy, hormone therapy and photodynamic therapy) according to PanDrugsdb (Piñeiro-Yañez et al, 2018). The drugs that remained unclassified were manually annotated. Finally, we computed the Pearson correlation between the viability change and the median regressed BCS for each drug category.

For Ho *et al*., we regressed the number of detected genes and the scores of G2M and S cell cycle phases. Five distinct TCs were found at resolution 0.2. The *bcSignatures* function was used to visualise the expression of BRAF inhibitor resistant biomarkers. Moreover, we generated a heatmap using the residuals of the regressed BCS to understand the differential behaviour of the therapeutic clusters using a selection of RAF and MEK inhibitors. Finally, we used the function *bc4Squares* to identify specific drugs to target TC5, BRAFi-resistant and parental cells.

In the case of the Kinker *et al*. study, we also obtained the BCS for the Recurrent Heterogeneous Programs (RHPs) described in the paper. Before moving on with the downstream analysis, we interrogated our clusters using the metadata information available. We noticed the cells primarily clustered based on the number of detected genes, the cell cycle phase and pool ID. Therefore, we proceeded to regress out these unwanted sources of variation and to recalculate the TCs at a resolution of 0.2. Nearest k neighbors was set to the default value of 20. As a result, 5 therapeutic clusters were identified. Finally, we computed the BCS statistics for both the individual TCs and cancer types and prioritised drugs according to their residual value and switch point. A heatmap (**Fig. 3a**) was generated using all cancer and cluster-specific drugs with a described mechanism of action and a heterogenous sensitivity pattern (corresponding to an SP filter between 0.4 and 0.6). In the heatmap, cells (columns) were first ordered by cluster, then by cancer type within each cluster. A hierarchical clustering was applied to the rows. On the other hand, the BCS matrix was correlated with the RHPs (**Fig. 3b**) and a total of 267 drugs were selected, corresponding to those with an absolute correlation higher than 0.6. The differential drug sensitivity analysis was performed using *limma* (Ritchie et al., 2015) on the PRISM repurposing dataset (Corsello et al. 2020). Specifically, we used log2 fold change data at a 2.5 µM dose from the secondary screen after replicate collapse. Annotations of the mechanisms of action were taken from the CLUE data library (clue.io/data). Lastly, a differential expression analysis between cells in TC2 and the rest of the TCs, was obtained using Seurat's *FindMarkers* function. The analysis showed an enrichment in the upregulation of EMT genes **(Fig. S13)**.

For the Jerby-Arnon *et al*. patient samples, the BCS were computed for each of the drugs featured in the $SS_C$ drug collection and for the functional immune resistance signature generated by the authors. For the immune resistance signature, we used Beyondcell's *GenerateGenesets* function to create a single bidirectional signature from the 44 genes comprising the up-regulated gene set and 69 genes in the down-regulated gene set. Beyondcell UMAP projection was calculated on the first 15 principal components of the BCS, as suggested by an elbow plot of the principal components against the explained standard deviation. Cluster resolution was set at 0.2 with 4 as nearest k neighbors instead of the default 20, due to the low number of sequenced cells (1881). Both the number of expressed genes and the scores for the S and G2M phases were regressed out.

#### Computation time

We benchmarked Beyondcell on an HP Z820 workstation featuring an Intel® Xeon® CPU E5-2670@2.60GHz by running 15 instances of the same analysis and recording the average time to completion, memory usage and CPU load. A regular analysis of a single-cell assay comprising 530 cells takes an average of 110 seconds and 4GB of RAM on a single thread to compute drug sensitivity scores, find therapeutic clusters and regress unwanted sources of variation such as the number of expressed genes or the cell cycle.

171(6), 1437-1452.
